# Requirement for specific bacterial genome maintenance pathways in repair of C8-linked pyrrolobenzodiazepine (PBD) bi-aryl monomer-mediated DNA damage

**DOI:** 10.1101/2022.06.10.495655

**Authors:** Asha Mary Joseph, Kazi Nahar, Saheli Daw, Md. Mahbub Hasan, Rebecca Lo, Tung B. K. Le, Khondaker Miraz Rahman, Anjana Badrinarayanan

## Abstract

Pyrrolobenzodiazepines (PBDs) are naturally occurring DNA binding compounds that possess anti-tumor and anti-bacterial activity. Chemical modifications of PBDs can result in improved DNA binding, sequence specificity and enhanced efficacy. More recently, synthetic PBD monomers have shown promise as payloads for antibody drug conjugates and antibacterial agents. The precise mechanism of action of these PBD monomers and their role in causing DNA damage remains to be elucidated. Here we characterized the damage-inducing potential of two C8-linked PBD bi-aryl monomers in *Caulobacter crescentus* and investigated the strategies employed by cells to repair the same. We show that these compounds cause DNA damage and efficiently kill bacteria, in a manner comparable to the extensively used DNA cross-linking agent mitomycin-C (MMC). However, in stark contrast to MMC which employs a mutagenic lesion tolerance pathway, we implicate essential functions for error-free mechanisms in repairing PBD monomer-mediated damage. We find that survival is severely compromised in cells lacking nucleotide excision repair and to a lesser extent, in cells with impaired recombination-based repair. Loss of nucleotide excision repair leads to significant increase in double-strand breaks, underscoring the critical role of this pathway in mediating repair of PBD-induced DNA lesions. Together, our study provides comprehensive insights into how mono-alkylating DNA-targeting therapeutic compounds like PBD monomers challenge cell growth, and identifies the specific mechanisms employed by the cell to counter the same.

## Introduction

Persistent DNA damage can be problematic to cells across domains of life, from unicellular bacteria to multicellular eukaryotes. It can have deleterious effects on basic cellular processes as well as organismal functions, and subsequently lead to cell death (Surova & Zhivotovsky, 2013). However, when designed and used appropriately, DNA damage can also work as tools to eliminate hazardous microorganisms or malignant tissues (de Almeida et al., 2021).

DNA damage can be mediated by endogenous or exogenous agents, leading to broad spectrum of DNA modifications (Chatterjee & Walker, 2017). For example, methlymethansulfonate (MMS) leads to methylation on a single base (G, A or C) causing mono-alkylated adducts such as 7-MeG, 1-MeA and 3-MeC (Beranek, 1990). UV exposure induces covalent linkages between adjacent pyrimidines, creating intra-strand crosslinks including cyclobutane pyrimidine dimers (CPDs) and pyrimidine-(6-4)-pyrimidone photoproducts (Chatterjee & Walker, 2017). On the contrary, mitomycin-C (MMC) causes inter-strand crosslinks in addition to mono-alkylated adducts and intra-strand crosslinks (Bargonetti et al., 2010; Tomasz, 1995).

Pyrrolobenzodiazepines (PBDs) are a group of DNA damaging agents that bind to the minor groove of DNA and alkylate DNA in a sequence-specific manner (Gerratana, 2012; Mantaj et al., 2017). PBDs typically contain an aromatic A-ring, a diazepine B-ring and a pyrrolidone C-ring, a structure that fits into the minor groove of DNA. Once secured within the minor grove, an electrophilic imine moiety in the B-ring establishes a covalent link with the C2-NH_2_ group of guanine base on the DNA, preferably within Pu-G-Pu sequences. Several PBDs including anthramycin, tomaymycin and sibiromycin, naturally produced by actinomycetes are mono-alkylators of DNA and exhibit strong anti-microbial and anti-tumor properties (Gerratana, 2012; Hurley, 1977; Leimgruber et al., 1965)

Diversity in PBDs can be typically brought about by the variations in the A- and C-rings (Mantaj et al., 2017). For instance, the A ring can be functionalized with electron-donating sidechains at the C7- and C8-positions resulting in increased alkylating potential and DNA binding capability of PBDs (Mantaj et al., 2017; Thurston et al., 1999). Research in the last three decades has led to development of several synthetic PBDs by engineering modifications to the basic PBD structure resulting in enhanced sequence specificity, stability and functionality (Bose et al., 1992; Gregson et al., 2001; Mantaj et al., 2017; Rahman et al., 2012).

Of these, synthetic PBD dimers formed by linking two PBD monomers via their aromatic A-ring has the potential to form cross-links on DNA, similar to cross-links formed by mitomycin C (MMC) and have been extensively studied for their chemotherapeutic value (Kung Sutherland et al., 2013; Puzanov et al., 2011; Rahman et al., 2011). Few of these dimers are being evaluated as payloads for antibody drug conjugates (ADCs) in clinical trials to specifically target and kill tumor cells (Kung Sutherland et al., 2013; Morgensztern et al., 2019). However, there have been a number of failures in the clinical development of PBD-dimer containing ADCs due to their toxicity (Jackson et al., 2018). This has resulted in significant interest in PBD-pseudodimer containing only one N10-C11 imine group, PBD-monomers and structurally related pyridinobenzodiazepines (PDD) monomers (Hoffmann et al., 2020; Kovtun et al., 2018), which are considered less toxic as they can only mono-alkylate DNA. C8-linked PBD bi-aryl monomers are a group of synthetic PBD monomers where a heteroaromatic group like pyrrole or imidazole is directly linked to a phenyl group using C-C coupling or fused with a phenyl ring and these bi-aryl units are attached to the C8 position of the PBD A-ring. These PBD monomers exhibit a strong preference for GC-rich DNA, leading to formation of mono-alkylated adducts on guanine. One such monomer, KMR-28-39 has shown low nanomolar to picomolar *in vitro* cytotoxicity against a panel of cancer cell lines and *in vivo* anti-tumor activity to breast cancer and pancreatic cancer xenografts in mouse models (Rahman et al., 2013). Interestingly, many of these PBD monomers also exhibited anti-bacterial activity against a range of Gram-positive bacteria, including methicillin-resistant Staphylococcus (Rahman et al., 2012). Another set of PBDs containing a terminal heteroaliphatic ring has similarly shown excellent activity against a panel of multidrug resistant Gram-negative bacteria (Picconi et al., 2020), without noticeable toxicity against eukaryotic cells. The therapeutic potential exhibited by PBD bi-aryl monomers (Andriollo et al., 2018; Brucoli et al., 2016; Picconi et al., 2020; Rahman et al., 2012; Rosado et al., 2011) is encouraging as they appear to possess significant anticancer and antibacterial activity and their eukaryotic toxicity can be tuned by altering the C8-side chain to make them selective against either eukaryotic or prokaryotic cells.

While induction of cross-links and DSBs by PBD dimers (Arnould et al., 2006; Jenkins et al., 1994) and their repair via endonuclease ERCC1 and homologous recombination (Hartley et al., 2010; Xing et al., 2019; Zhong et al., 2019) are well characterized, there has been little research in elucidating the mechanism of action of PBD monomers beyond their ability to inhibit transcription factors (Corcoran et al., 2019; Kotecha et al., 2008; Rahman et al., 2013). The contribution of DNA-damage mediated effect on their overall cytotoxicity or antibacterial activity needs to be studied to properly evaluate their potential as chemotherapeutic agents. Furthermore, as efforts progress towards developing DNA-interactive PBD monomers as payloads for antibody drug conjugate, it is also important to elucidate the mechanism(s) that lead to repair of such PBD lesions, and identify the outcome of their repair on both survival and mutagenesis.

The main objective of this work was to identify the pathways(s) involved in the repair or tolerance of lesions induced by PBD monomers and assess the possible involvement of error-prone repair or tolerance mechanisms (such as translesion synthesis) that can impact damage-induced mutagenesis, contributing to development of resistance. We used the non-pathogenic Gram-negative bacteria *Caulobacter crescentus* as our model system. Unlike *E. coli*, this GC-rich organism shares several key genome maintenance features, including error-prone lesion tolerance mechanisms, with pathogenic bacteria such as *Pseudomonas aeruginosa* and *Mycobacterium tuberculosis* (Alves et al., 2017; Boshoff et al., 2003; Galhardo, 2005; Jatsenko et al., 2017; Warner et al., 2010). Using two prototypes (described in Fig. 1A and below), we found that C8-linked PBD bi-aryl monomers induced DNA damage and efficiently killed *Caulobacter crescentus*. Repair of these lesions was predominantly mediated by nucleotide excision repair; lack of repair led to generation of double-strand breaks and severely compromised survival. In contrast to MMC we found that mutagenic translesion synthesis was not essential for PBD monomer-mediated damage tolerance or repair. Taken together, our study uncovers, for the first time, the mechanisms involved in repair of DNA-monoalkylations induced by PBD monomers and their overall impact on genome integrity and survival of bacterial cells.

**Figure 1:**
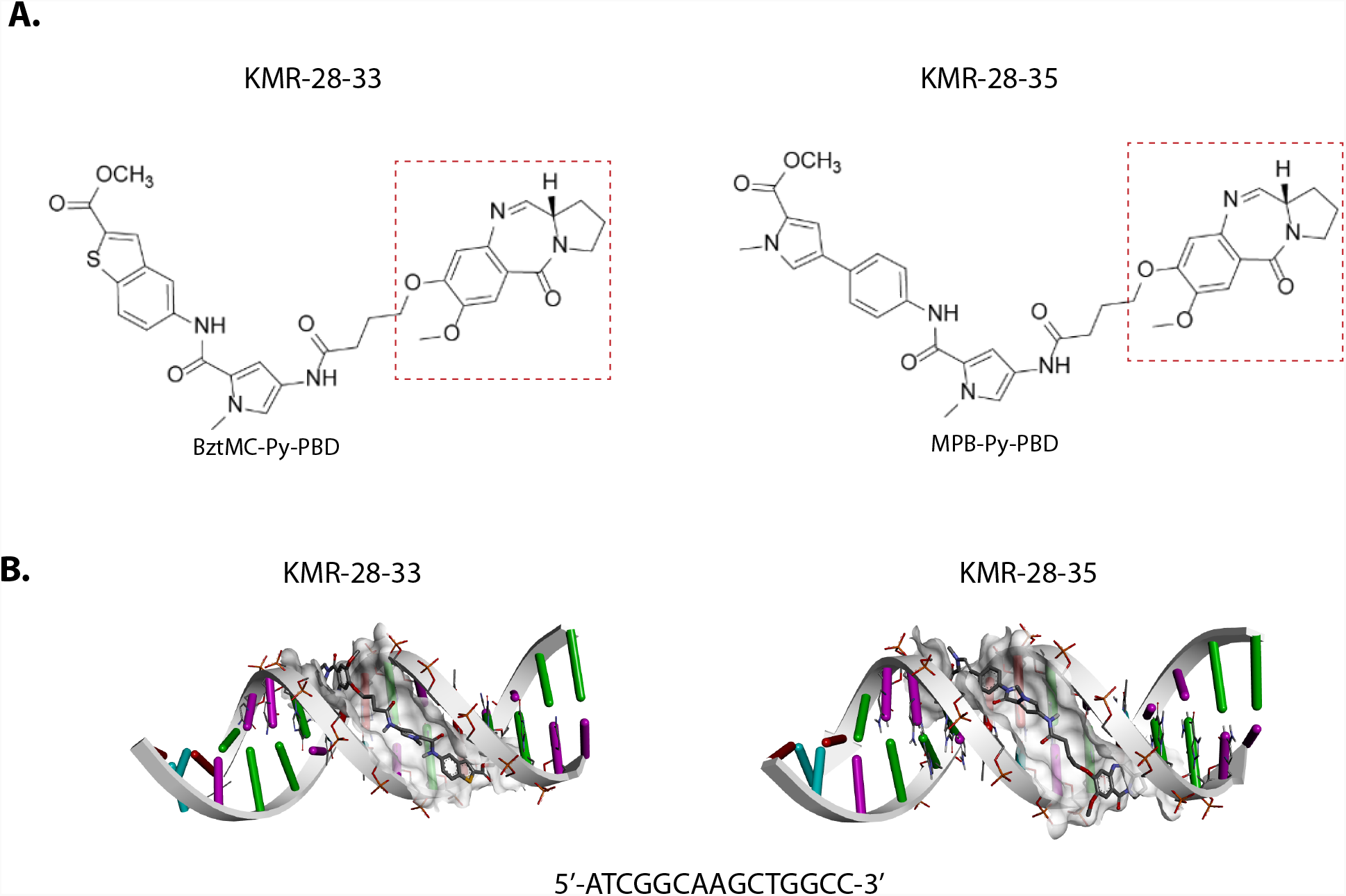
C8-linked PBD bi-aryl monomers KMR-28-33 and KMR-28-35. (A): Structures of KMR-28-33 and KMR-28-35; pyrrolo(2,1-c)(1,4)benzodiazepine (PBD) unit is highlighted by the red square, Py - 1-methylpyrrol-3-amine, BztMC - methyl 5-aminobenzothiophene-2-carboxylate, MPB - 4-(1-methyl-1H-pyrrol-3-yl)benzenamine. (B) Molecular docking of KMR-28-33 and KMR-28-35 with 15 bp DNA sequence taken from the ORF of *dnaE* gene (5’-ATCGGCAAGCTGGCC-3’; GC content – 66%) suggests snug fit of both KMR-28-33 and KMR-28-35 within the DNA minor groove.

## Results

### Chemically synthesized C8-linked PBD bi-aryl monomers KMR-28-33 and KMR-28-35 cause DNA damage in *Caulobacter crescentus*

To assess the DNA damaging potential of PBD monomers, we chose two C8-linked PBD bi-aryl monomers (KMR-28-33 and KMR-28-35) (Fig. 1A) as prototypes. These compounds, which have propensity to bind to DNA (preferentially to GC-rich tracts (Rahman et al., 2013)), showed strong cytotoxic (Rahman et al., 2013) and antibacterial activity towards Gram-positive bacteria in the earlier studies (Rahman et al., 2012). As a comparison, we evaluated the repair of lesions caused by a conventional and well-characterized DNA cross-linking agent (mitomycin-C, MMC).

Indeed, molecular modelling of KMR-28-33 and KMR-28-35 binding to DNA also lent strong support to the possibility that these PBD monomers could result in DNA lesions, without distorting the DNA helix itself, as they snugly fit within the DNA minor groove (Fig. 1B, S2A. S2B). Our *in vitro* FRET-based DNA melting assays corroborated the binding of these PBD monomers to DNA (Fig. S2C). Given these observations, we investigate whether these PBD monomers cause DNA damage and if so, what mechanisms are employed by cells to repair the same. For this, we use the GC-rich model system *Caulobacter crescentus*, where we deleted genes involved in repair of specific types of DNA damage (Fig. S2E).

We first assessed cell survival upon treatment with KMR-28-33 and KMR-28-35 (synthesis described in Fig. S1A-B) in a wild type background. As a reference, we also exposed cells to well-characterized DNA damaging agents, that are known to induce specific types of damage (Fig. 2A and S2D). In particular, we compared the effects of KMR-28-33 and KMR-28-35 to mitomycin-C (MMC). Both KMR-28-33 and KMR-28-35 can only form DNA monoadducts while MMC can form intra-strand and inter-strand crosslinks in addition to monoadducts (Rahman et al., 2012; Warren et al., 1998).

**Figure 2:**
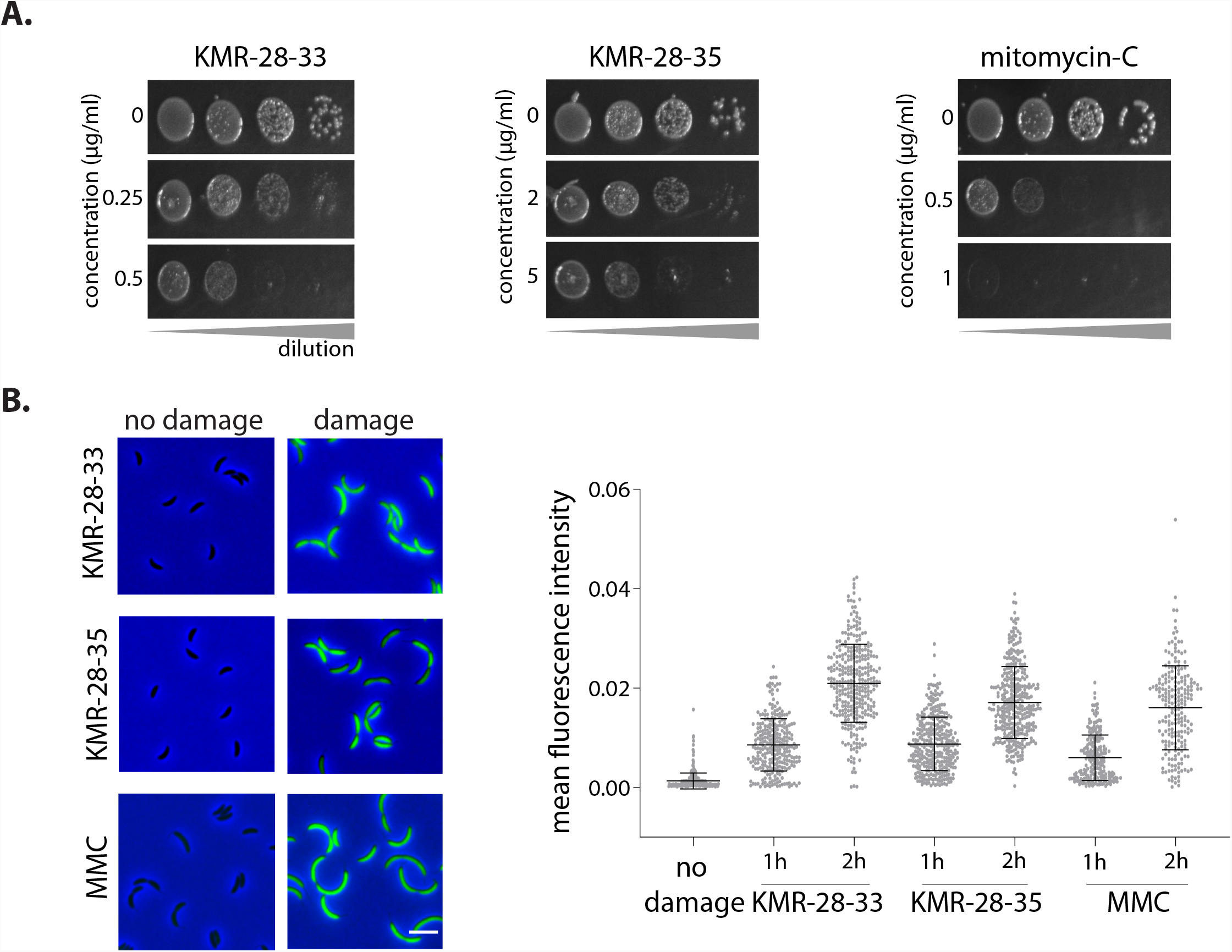
KMR-28-33 and KMR-28-35 treatment causes cell death and induces the SOS response. (A) Representative images of wild type *Caulobacter crescentus* growth on increasing concentrations of DNA damaging agents KMR-28-33, KMR-28-35 and MMC. Grey triangle at the bottom of each image panel depicts increasing dilution of the bacterial culture from left to right. Minimum of two independent experiments were performed for each dose. (B) [left] SOS induction is measured by assessing the expression of YFP from an SOS-inducible promoter (*P*_*sidA*_*-yfp*). Representative images of cells expressing the reporter with and without treatment with KMR-28-33 (0.5 µg/ml), KMR-28-35 (1 µg/ml) or MMC (0.5 µg/ml). Scale bar – 4 µm. [right] Total fluorescence intensity normalized to cell area plotted for indicated duration of damage treatment. Each dot represents a single cell. Mean and SD are shown in black (n ≥ 215).

We observed that both C8-linked PBD bi-aryl monomers caused cell death in a dose-dependent manner (Fig. 2A). We next asked whether these PBD monomers resulted in DNA damage. For this, we measured SOS response induction in wild type *Caulobacter* after treatment with the KMR-28-33 and KMR-28-35. We specifically quantified the expression of a fluorescence marker (*YFP*) induced under an SOS promoter (P_*sidA*_) integrated on the *Caulobacter* chromosome at the *xyl* locus (Badrinarayanan et al., 2015; Joseph et al., 2021). We found that exposure of cells to both PBD monomers resulted in significant accumulation of SOS-induced YFP (comparable to that observed in case of MMC, at concentrations that similarly affected cell growth in all three damages), suggesting that KMR-28-33 and KMR-28-35 caused DNA damage (Fig. 2B).

### Requirement for RecA, but not the SOS response, in C8-linked PBD bi-aryl monomer-treated cells

Based on the above observations we wondered whether specific pathways under the SOS response were required to repair KMR-28-33 and KMR-28-35-mediated damage. For example, in the case of MMC, the error-prone translesion synthesis polymerase, DnaE2, has been found to be essential (Boshoff et al., 2003; Galhardo, 2005; Joseph et al., 2021). In support of the possibility that these PBD monomers indeed induce DNA damage, we found that cells lacking *recA* were compromised for survival upon treatment with both PBD-monomers (Fig. 3A). The difference in sensitivity for Δ*recA* cells across KMR-28-33, KMR-38-35 and MMC suggested that there may be distinct repair mechanisms at play. We thus uncoupled key DNA damage-specific functions of RecA (recombination and SOS induction), to assess the contribution of the two towards survival. For this, we generated a strain where the SOS repressor, *lexA*, is deleted. To circumvent the problem of cell length elongation in this background, previous studies have additionally deleted the SOS-induced division inhibitor, *sidA* (Modell et al., 2011). In this constitutive ‘SOS-ON’ background, deletion of *recA* would predominantly eliminate its function in recombination. Thus, this triple deletion of *lexAsidArecA* can be used as a genetic read-out to test requirement of SOS vs recombination functions of RecA upon DNA damage treatment.

**Figure 3:**
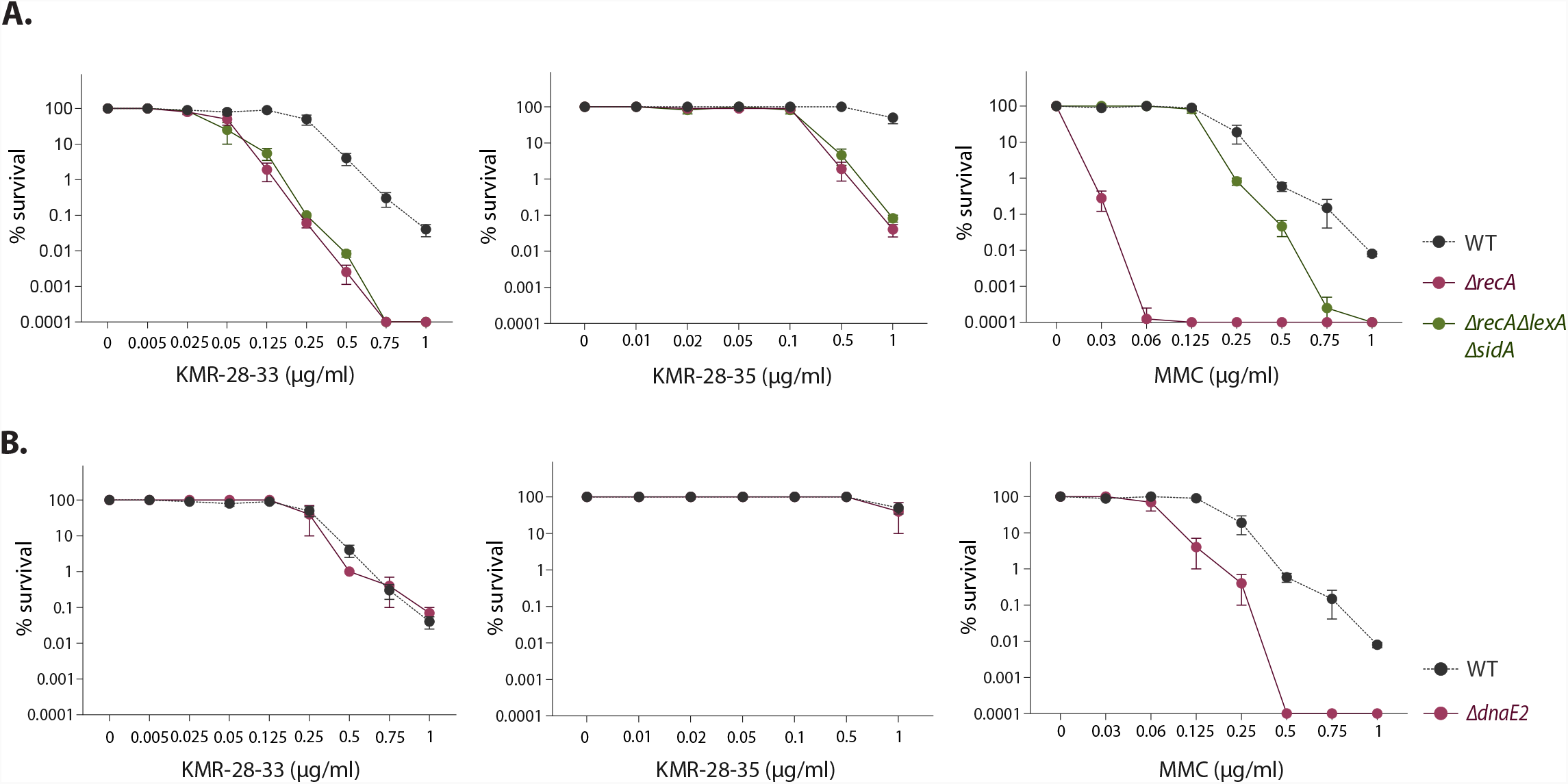
Requirement for RecA, but not the SOS response, in C8-linked PBD bi-aryl monomer-treated cells. (A) Survival of wild type, *ΔrecA* and *ΔrecAΔlexAΔsidA* strains under increasing doses of KMR-28-33, KMR-28-35 and MMC. Minimum of three independent experiments were performed for each strain. Mean and SEM from all repeats for each strain is plotted. (B) Survival of wild type and *ΔdnaE2* strains under increasing doses of KMR-28-33, KMR-28-35 and MMC. Minimum of three independent experiments were performed for each strain. Mean and SEM from all repeats for each strain is plotted (wild type data from Figure 3A for comparison).

In case of MMC-damage, SOS-ON cells performed better than those deleted for *recA* (Fig. 3A), suggesting that a pathway regulated under the SOS response contributed significantly to cell survival under MMC damage. Indeed, this phenotype can be attributed to the expression of the TLS pathway (including the error-prone polymerase DnaE2) in SOS-ON cells, but not in cells deleted for *recA*. Previous studies have implicated an important role for this mechanism in tolerance of MMC-induced lesions, independent of *recA* (Galhardo, 2005; Joseph et al., 2021) (Fig. 3B). In contrast to MMC, we found that Δ*recA* or Δ*lexAsidArecA* cells were similarly compromised in growth when treated with the C8-linked PBD bi-aryl monomers, suggesting that SOS function was not required for combatting KMR-28-33 and KMR-28-35-mediated damage (Fig. 3A). In line with this observation, we also found that TLS polymerase DnaE2 was not essential to tolerate C8-linked PBD bi-aryl monomer-mediated damage (Fig. 3B).

Given the sensitivity of Δ*recA* cells (independent of SOS) to treatment with the KMR compounds, we asked whether recombination-mediated repair contributed to cell survival under PBD monomer-mediated damage. For this we deleted genes involved in specific recombination-based repair: a. *recF, recO* and *recR* that function in single-strand gap (SSG) repair and b. *addAB* and *recN* that function in double-strand break (DSB) repair (Rocha et al., 2005; Spies & Kowalczykowski, 2014). It is important to note that although we categorize the genes in this manner, there is also evidence to suggest that they may have overlapping functions (Pages, 2003).

In case of MMC damage, we found that cells deleted for *recF, recO* or *recR* were similarly sensitive to damage, and comparable with a *recA* deletion (Fig. 4A-B). Given the Δ*recA*-like sensitivity in these backgrounds, it is tempting to speculate that these proteins may play a role in loading RecA at ssDNA gaps to enable SOS induction, apart from contribution to recombination-based repair. In line with this possibility, *addAB* and *recN* deleted cells were less compromised in growth when compared to the Δ*recA* cells (Fig. 4A). In contrast to MMC damage, cells treated with KMR-28-33 or KMR-28-35 were similarly compromised in growth in the absence of SSG or DSB repair (Fig. 4A-B). Importantly, the sensitivity observed in case of these deletions mirrored that of *recA*, suggesting that both recombination pathways may contribute to repairing PBD bi-aryl monomer-induced damage.

**Figure 4:**
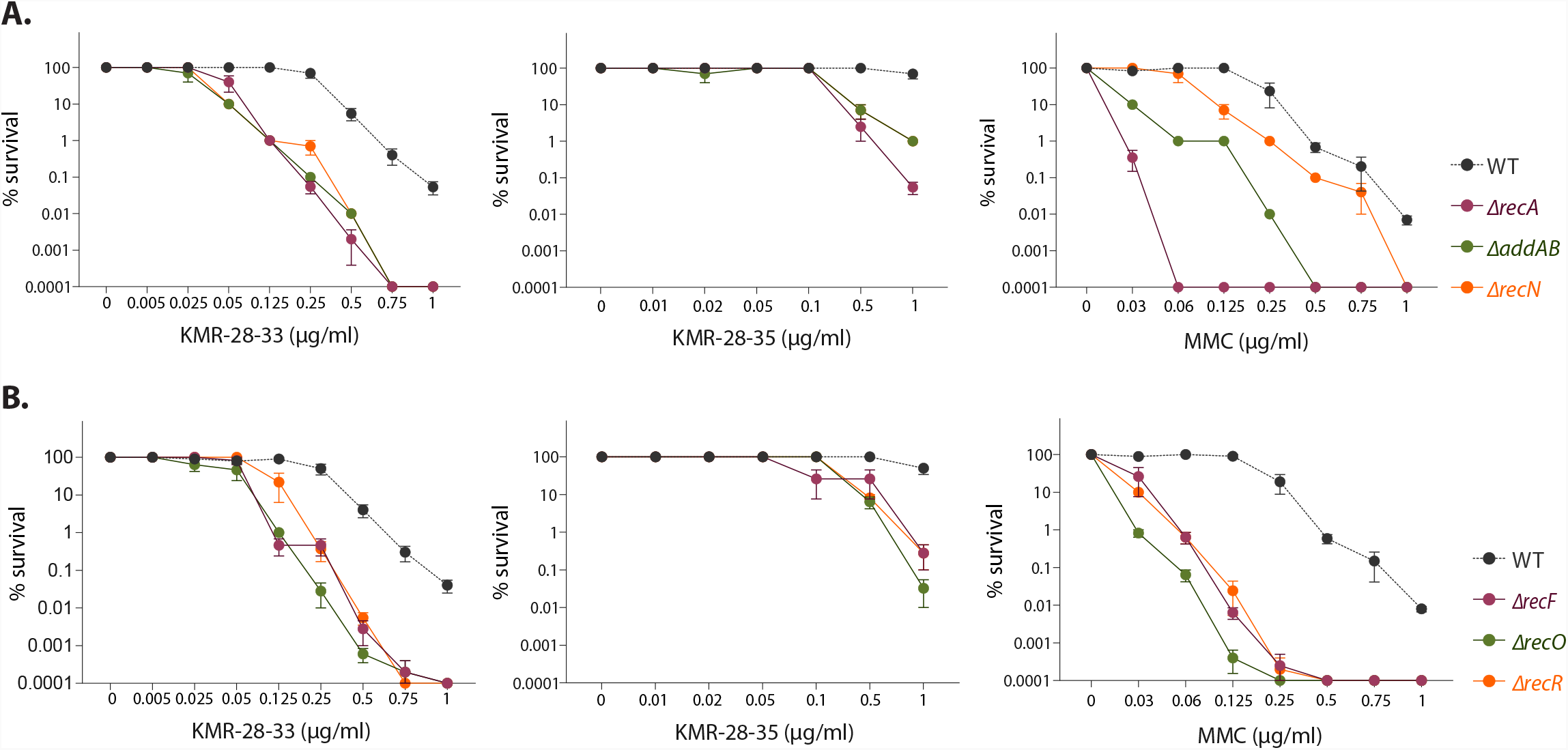
Recombination-mediated repair contributes to survival under DNA damage caused by KMR-28-33 and KMR-28-35. (A) Survival of wild type, *ΔrecA, ΔaddAB* and *ΔrecN* strains under increasing doses of KMR-28-33, KMR-28-35 and MMC. Minimum of three independent experiments were performed for each strain. Mean and SEM from all repeats for each strain is plotted (wild type and *ΔrecA* data from Figure 3A for comparison). (B) Survival of wild type, *ΔrecF, ΔrecO* and *ΔrecR* strains under increasing doses of KMR-28-33, KMR-28-35 and MMC. Minimum of three independent experiments were performed for each strain. Mean and SEM from all repeats for each strain is plotted (wild type data from Figure 3A for comparison).

### Nucleotide excision repair (NER) is essential for survival under KMR-28-33 and KMR-28-35-induced DNA damage

Although we implicated a role for recombination in repair of C8-linked PBD bi-aryl monomer-mediated damage, these PBD monomers are thought to predominantly form DNA mono-alkylations and are not known to directly induce double-strand breaks. To estimate the DSB-inducing potential of KMR-28-33 and KMR-28-35, we adapted the Gam-GFP reporter system previously described in *E. coli* (Shee et al., 2013) to mark DSB ends *in vivo* in *Caulobacter*. Using this system, we estimated the percentage of cells with Gam localization in the presence and absence of the lesion-inducing damaging agents (Fig. S3A). In the absence of any damage, <1% cells had detectable foci. As anticipated, in the presence of the KMR compounds, this number increased only nominally to ∼5% after 2 h of treatment with the compounds. This was similar to observations made for MMC-induced damage as well (Fig. S3A). Interestingly loss of *recN*, required for recombination repair, only resulted in a modest increase in DSBs under KMR compounds or MMC (Fig. S3A). Together, this suggested to us that recombination may only be a minor repair pathway, with some other mechanism(s) likely enabling lesion repair or tolerance.

We thus wondered which lesion repair or tolerance pathways were required to repair damage induced by the C8-linked PBD bi-aryl monomers. As shown earlier, unlike MMC, we had ruled out a role for TLS polymerase DnaE2 (Fig. 3A-3B). Indeed, the lack of SOS response essentiality also eliminated a role for the other TLS polymerase, DinB (Galhardo, 2005; Joseph & Badrinarayanan, 2020), in this case (Fig. 3A). We next assessed survival in cells compromised for alkylation repair (*alkB*) (Colombi & Gomes, 1997) or mismatch repair (*mutL*) (Chai et al., 2021) and found that these pathways also did not contribute to repair of KMR-induced damage (Fig. S3B).

We thus turned to Nucleotide Excision Repair (NER). NER is an important mechanism of DNA lesion repair, that predominantly acts on helix-distorting lesions (Jia et al., 2009; Liu et al., 2011). Although damage induced by these PBD monomers is thought to not cause significant distortion to the DNA helix, we found that cells lacking *uvrA* (the lesion scanning component of the NER pathway) were severely compromised in survival under KMR-28-33 or KMR-28-35 damage (Fig. 5A). This was in contrast to MMC, where UvrA is required but not as essential, as seen in case of the C8-linked PBD bi-aryl monomers (Fig. 5A). Indeed, differential sensitivity of Δ*uvrA* strains to UV, MMS and norfloxacin damage further underscored the specificity of this repair pathway (Fig. S4A).

**Figure 5:**
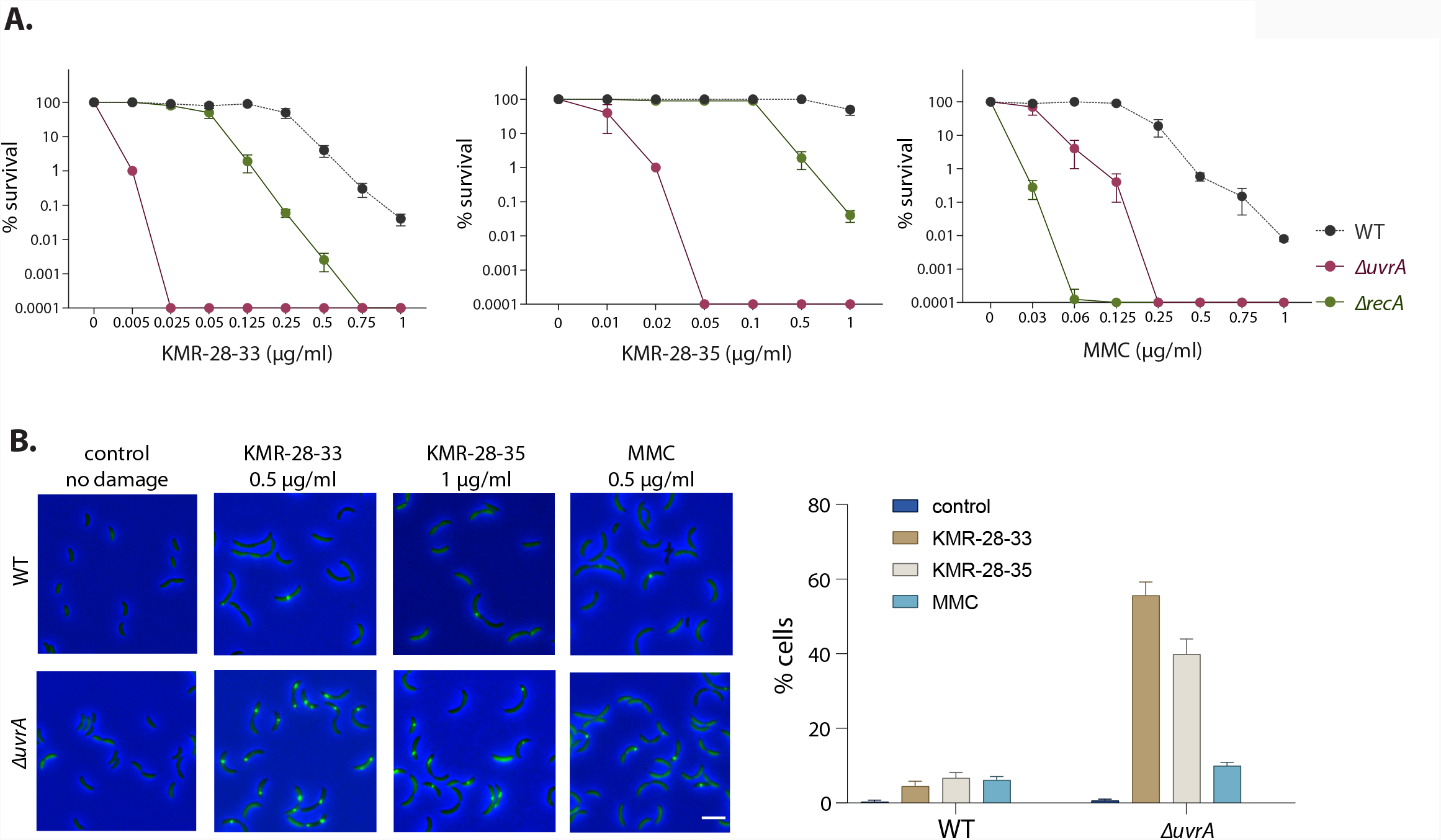
Nucleotide excision repair (NER) is essential for survival under KMR-28-33 and KMR-28-35-induced DNA damage. (A) Survival of wild type, *ΔrecA* and *ΔuvrA* strains under increasing doses of KMR-28-33, KMR-28-35 and MMC. Minimum of three independent experiments were performed for each strain. Mean and SEM from all repeats for each strain was plotted (wild type and *ΔrecA* data from Figure 3A for comparison). (B) [left] Representative images for cells with Gam-GFP foci upon treatment with KMR-28_33, KMR-28-35 and MMC in wild type and *ΔuvrA* strains (wild type images from Figure S3A for comparison). Scale bar – 4 µm. [right] Percentage cells with Gam-GFP foci upon treatment with KMR-28-33, KMR-28-35 and MMC in wild type and *ΔuvrA* strains (wild type data from Figure S3A for comparison). Mean and SD for data from three independent experiments is plotted (n ≥ 330 cells).

NER primarily functions via two ways: transcription-coupled repair (TCR) and global genomic repair (GGR). In case of TCR, Mfd plays a central role in recruiting Uvr components to the site of lesion for excision, followed by gap filling (C. Selby & Sancar, 1993; Strick & Portman, 2019). On the other hand, in case of GGR, UvrA is thought to scan and recognize lesions across the genome, and subsequently initiate repair (Kisker et al., 2013). We thus deleted the *mfd* homolog in *Caulobacter* and found that cells lacking the ability to engage in TCR were not as severely compromised in survival, when compared to Δ*uvrA* deleted cells (Fig. S4B). These data suggest that the GGR arm of NER is primarily responsible for repair of DNA lesions induced by KMR-28-33 and KMR-28-35. Indeed, alternate pathways for repair coupled to transcription, independent of Mfd, have also been proposed (C. P. Selby, 2017). We have not investigated the role of such mechanisms, which are currently not well-characterized or identified in *Caulobacter*.

Importantly, the absence of NER resulted in severe genome instability in cells treated with KMR-28-33 or KMR-28-35. As shown above, only 5-6% wild type cells treated with KMR compounds or with MMC had Gam localizations, indicating DSBs. In contrast, the lack of *uvrA* resulted in a significant increase in cells with Gam localizations in case of the PBD monomers, with 56% cells having DSBs upon KMR-28-33 treatment and 40% cells with DSBs on KMR-28-35 treatment (Fig. 5B). This was not found to be the case for MMC-treated cells, where *uvrA* deletion only led to modest increase in percentage cells with DSBs (6% in wild type to 10% in NER-compromised cells) (Fig. 5B). Such marked genome instability was observed only when NER action was compromised, as deletion of *recN* did not result in increase in localization of Gam-GFP (Fig. S3A).

Together these results highlight: a. the essentiality of NER in repairing C8-linked PBD bi-aryl monomer-mediated damage and b. the distinct mechanisms of DNA damage repair in case of the PBD monomers, when compared to an extensively studied DNA cross-linking agent, MMC (Fig. S5).

## Discussion

Earlier studies have affirmed the specificity of action of pathways for repair of DNA damage. For example, MrfA and MrfB in *Bacillus subtilis*, and MmcA and MmcB in *Caulobacter crescentus* are essential for repair of only MMC-induced lesions (Burby & Simmons, 2019; Lopes-Kulishev et al., 2015). Similarly, NER has been implicated as the primary repair pathway in case of nitrofurazone damage in *E. coli* (Ona et al., 2009). In many instances, the requirement for different repair components is likely driven the structural variations in lesions induced by specific damaging agents (Cole et al., 2018; Ona et al., 2009; Williams et al., 2013). Such difference in function can be observed even within a pathway, when the type of lesion differs. Both prokaryotic and eukaryotic TLS polymerases exhibit substrate specificity which defines their lesion bypass properties including efficiency of bypass and fidelity (Inomata et al., 2021; Ippoliti et al., 2012; Prakash et al., 2005; Waters et al., 2009). In case of *Caulobacter*, requirement for TLS polymerase DnaE2 is relatively higher for repair of MMC lesions than UV lesions (Galhardo, 2005; Joseph et al., 2021).

Harnessing this specificity in function of repair pathways, in this study, we determined the damage-inducing potential of two C8-linked PBD bi-aryl monomers and delineated the strategies employed by bacterial cells to repair the same. Our results indicate that base modifications (in the form of mono-alkylated adducts) caused by the KMR compounds are predominantly repaired by NER and do not employ error-prone TLS mechanisms. Indeed, it is tempting to attribute the difference between MMC and the KMR compounds to repair/ tolerance of monoalkylations vs inter and intra-strand DNA crosslinks. While the C8-linked PBD bi-aryl monomers can only form mono-adducts, MMC treatment can result in mono-adducts as well as both inter and intra-strand DNA crosslinks. Thus, the relative contribution of various repair components could differ between lesions that are structurally and chemically distinct, but mechanistically similar.

In addition to NER, we find contribution of recombination-mediated repair to cell survival in case of the KMR compounds. The exact sequence of event(s) that lead to conversion of a mono-adduct into a DSB remains elusive. It is speculated that cellular processes like transcription, replication and even incomplete repair can lead to generation of single stranded gaps as well as double stranded breaks (Aguilera & Gaillard, 2014; Mehta & Haber, 2014). The observation that the NER mutant is far more compromised in survival and generates much higher proportion of DSBs than a recombination mutant supports the idea that recombination may act secondary to lesion repair via NER. Indeed, we did consider making a strain impaired in both NER and recombination to test this possibility. However, the very high sensitivity of the Δ*uvrA* strain to KMR compounds precludes our ability to do so with reliability. Future work aimed at quantitative estimation of the levels and types of DNA damage induced *in vivo* in all three cases (KMR compounds and MMC) will enable us to further discern the hierarchy of requirement and action of repair pathways.

In sum, our work highlights the importance of studying the mechanism of action of potential DNA-interactive therapeutics like PBD monomers in depth, to understand how they may affect cell growth and what strategies may be employed by the cell to respond to the same. For example, when considering a DNA damaging agent for therapeutic purposes, it is important to understand the fidelity of repair mechanisms that could be employed by the cells. Mutagenic repair can be a major source of stress-induced mutagenesis and subsequent development of resistance (Fitzgerald et al., 2017; Ippoliti et al., 2012; Joseph & Badrinarayanan, 2020). Our findings suggest that C8-linked PBD bi-aryl monomer-induced lesions are likely non-mutagenic, and are predominantly repaired by nucleotide excision repair, thus negating an important driver for development of chemoresistance and antimicrobial resistance. This contrasts with PBD dimers which are known to cause mutagenesis and non-selective toxicity, and makes the case for using PBD monomers as ADC payloads to overcome recent clinical failures observed with PBD dimers (Jackson et al., 2018). Identifying the dependency on NER (specifically Uvr components, which are restricted to bacteria) further opens up possibilities for considering inhibitors for Uvr components to use in combination with the PBD monomers. Indeed, chemical inhibitors for specific repair pathways have been identified previously, including for *E. coli* RecBCD, *H. pylori* AddAB and *M. smegmatis* Uvr proteins (Amundsen et al., 2012; Mazloum et al., 2011). Combining a DNA damaging drug with a small molecule inhibitor capable of dampening damage repair in the pathogen can potentiate the efficacy of the drug and reduce pleotropic cytotoxicity (Lim et al., 2019).

## Materials and methods

### Synthesis of KMR-28-33 and KMR-28-35

The PBD component of the hybrids was synthesized from vanillin as previously described in the literature (Rahman et al., 2013) and is summarized in Fig. S1A. A four-carbon linker was used to connect the PBD component with the non-covalently interactive subunits, as chains of this length had proven optimal in previous hybrid SAR studies (Wells et al., 2006). The linker was located at the C8 position of the molecule to allow an isohelical fit of the non-covalent component of the hybrid along the minor groove upon covalent PBD binding. After the synthesis of the PBD core, non-covalently interactive side chains were constructed using combinations of benzofused (benzothiophene, KMR-28-33), five membered heterocyclic structures (N-methyl pyrrole/N-methyl imidazole) and MPB (4-(1-methyl-1H-pyrrol-3-yl)benzenamine, KMR-28-35) moieties. The MPB subunit was synthesized using Suzuki-Miyuara conditions previously described (Rahman et al., 2013) (Fig. S1B). These moieties were linked via Steglich amide bond formation at positions which maintained the overall fit of the hybrid for the DNA minor groove (C2/C5 for benzofused, C1/C4 for heterocyclic components), and finally the N10/C11 imine component of the molecule was activated using tetrakis palladium and pyrrolidine.

### Bacterial strains and growth conditions

Bacterial strains, plasmids and primers used in the study are listed in Table S1-S3. Chromosomal deletions and integrations were performed using either a two-step recombination method with a *sacB* counter-selection marker (Skerker et al., 2005) or using integrating vectors from Thanbichler et al. (Thanbichler et al., 2007). Transductions were carried out with ΦCR30 (Ely, 1991). *Caulobacter crescentus* cultures were routinely grown at 30°C in PYE (0.2% peptone, 0.1% yeast extract and 0.06% MgSO_4_). For strains expressing *Gam-GFP* under P_*xyl*_, 0.3% xylose was added 3h prior to imaging.

### Survival assay

*Caulobacter cultures* were grown in PYE to O.D_600_ of 0.3. Serial dilutions in 10-fold increments were made and 6 µl of each dilution (10^−1^ to 10^−8^) were spotted on PYE agar containing appropriate concentrations of different DNA damaging agents. For UV damage, serial dilutions of the culture were spotted on PYE agar plates and exposed to specific energy settings in a UV Stratalinker 1800 (STRATAGENE). Growth was assessed from the number of spots on the plates after two days of incubation at 30°C. Percentage survival for each strain was calculated by normalizing growth of that specific strain on different doses of DNA damage to that on media without DNA damage.

### Fluorescence microscopy and image analysis

Saturated overnight cultures were back diluted in fresh PYE and allowed to grow at least for two generations (approx. 3h) until OD_600_ was 0.1. Images were taken without damage treatment (no damage control) and after treatment with specified doses of DNA damage. 1 ml aliquots of cultures were taken at specified time points, pelleted and resuspended in 100 µl of growth medium. 2 µl of cell suspension was spotted on 1% agarose pads (prepared in water). Imaging was performed on a wide-field epifluorescence microscope (Eclipse Ti-2E, Nikon) equipped with a 60X oil immersion objective (plan apochromat objective with NA 1.41) and pE4000 light source (CoolLED). Images were acquired with Hamamatsu Orca Flash 4.0 camera using NIS-elements software (version 5.1). For quantifying YFP induction under *P*_*sidA*_ promoter, cells were segmented using Oufti (Paintdakhi et al., 2016) in MatLab, and florescence intensities normalized to cell lengths were extracted. Percentage cells with DSBs were quantified by counting cells with Gam-GFP foci using the Cell Counter plugin in ImageJ. Graphs were plotted in GraphPad Prism 7.

### Molecular Modelling

The 3D structures of desired B-form DNA sequences from the NA1000 (*Caulobacter crescentus*) genome sequence (15 bp from *dnaE* ORF: 5’-ATCGGCAAGCTGGCC-3’, LexA box within the promoter of *recA*: 5’-GTTCGCAAGATGTTC-3’ and *CCNA_RR0074 sRNA*: 5’-CCCCTTCGCCCTCCT-3’,) were generated using PyMOL 2.5 structure Builder. For small molecule ligands used in this study, 3D structures were generated using Chem3D 20.0 program. The DNA structures were processed (energy minimization and addition of polar hydrogens) using MGLTools v1.5.7 (https://autodock.scripps.edu/). The grid box was configured for each DNA macromolecule to cover the whole length of the structure so that the ligand was able to find best possible binding sites along with the DNA structures including both the major and minor grooves. The small molecular ligands were also processed with the same tools. Finally, the molecular docking was performed using opensource AutoDock Vina v.1.2.0 (https://vina.scripps.edu/) (Trott & Olson, 2009). The default flexible docking parameters were kept for docking. The post processing of the output files was curated using PyMOL 2.5 and the molecular interactions were visualized using BIOVIA Discovery Studio Visualizer.

### FRET-based DNA melting

All FRET duplexes and hairpins were purchased as pairs of complimentary or self-complimentary single-stranded oligonucleotides in lyophilised form from Eurogentec Ltd. The oligonucleotides were fluoro-tagged at the 5’ position with TAM and 3’ position with TAMRA. Sequences used were as follows; AT-rich sequence (seq-1): 5’-FAM-TAT-ATA-TAG-ATA-TTT-TTT-TAT-CTA-TAT-ATA-TAMRA-3’; GC-rich sequence (seq-2): 5’-FAM-TAT-AGG-GAC-AGC-CCT-ATA-3’, 3’-TAMRA-ATA-TCC-CTG-TCG-GGA-TAT-5’. Nuclease-free water was used to prepare stock solutions (20 μM) of the oligonucleotide hairpins/duplex strands. These stock solutions were diluted to concentrations of 400 nM using FRET buffer (50 mM potassium cacodylate, pH 7.4). The solutions were then heated to 85 °C/80 °C for five/ten minutes (hairpin/duplex solutions, respectively) using a heating block (Grant-Bio). The solutions were allowed to cool to room temperature overnight and cooled to -20 °C to complete the annealing process. Annealed stock solutions were diluted to concentrations of 100 and 10 nM using FRET buffer to prepare working solutions. PBD monomers, GWL-78 (Wells et al., 2006) and mitomycin C to be incubated with the DNA duplexes were dissolved in DMSO to form 5 mM solutions. Working solutions of PBD monomers and mitomycin C (5 μM and 1 μM) were prepared using FRET buffer. The working solutions of the compounds and DNA hairpins/duplexes were mixed (1:1 ratio, 25 μL of each solution) in the wells of a 96 well plate (Bio-Rad). The wells were covered and placed in a DNA Engine Opticon system for melting. The samples were heated over a range of 30-100 °C, with fluorescence readings (incident radiation 450-495 nm, detection 515-545 nm) taken at intervals of 0.5 °C. Experimental data was imported into Origin (OriginLab Corp.), where the curves were smoothed and normalised. Using a script, the point of inflection of the first derivative of the melting point for each sample on the plate was calculated. The difference between the melting temperature of each sample and that of the blank (i.e., the ΔTm) was used for comparative purposes. Mean is shown from three independent repeats.

## Author contributions

AJ led the project, generated tools and reagents, carried out *in vivo* experiments in *Caulobacter* and conducted data analysis. SD contributed tools and reagents, and carried out *in vivo* experiments. KN and MMH carried out experiments pertaining to the KMR compounds synthesis and *in vitro* characterization. RL and TL contributed tools and reagents. KMR and AB conceived and supervised the project. AJ, KMR and AB procured funding and wrote the manuscript, with feedback from all authors.

## Acknowledgements

AJ and AB thank members of the AB lab for feedback on the work. AJ acknowledges support from DST N-PDF SERB. AB acknowledges support from the DBT-IYBA grant and intra-mural funding from NCBS-TIFR. TL acknowledges support from the Royal Society University Research Fellowship Renewal (URF\R\201020) and BBSRC (BBS/E/J/000PR9791).

## Supplementary Information

### Supplementary Figures

**Figure S1:**
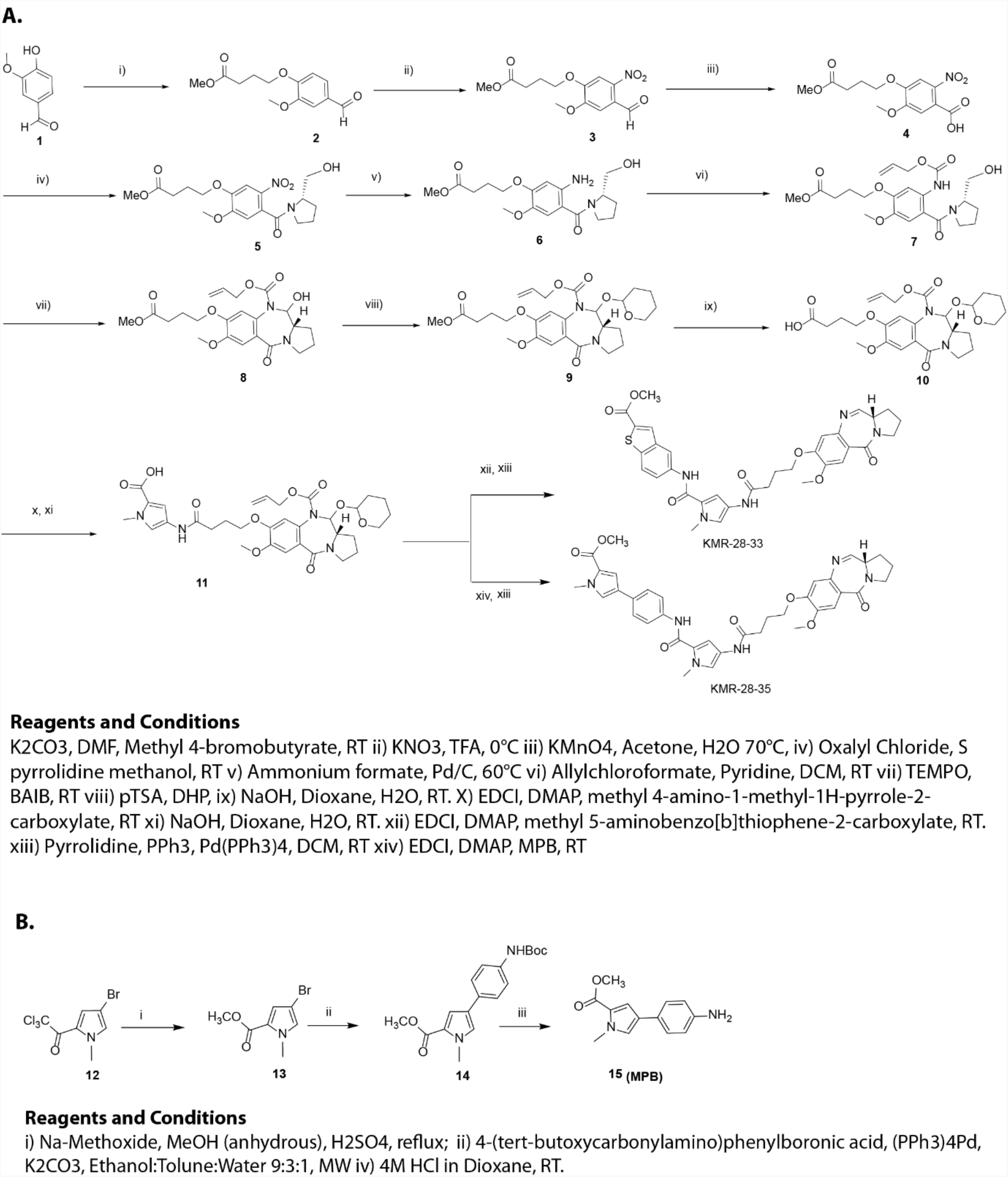
(A) General synthetic scheme for the synthesis of PBD core (10) and the conjugation of C8-side chain to the PBD core to obtain KMR-28-33 and KMR-28-35. (B) Synthesis of MPB building block 15 which is present in KMR-28-35.

**Figure S2:**
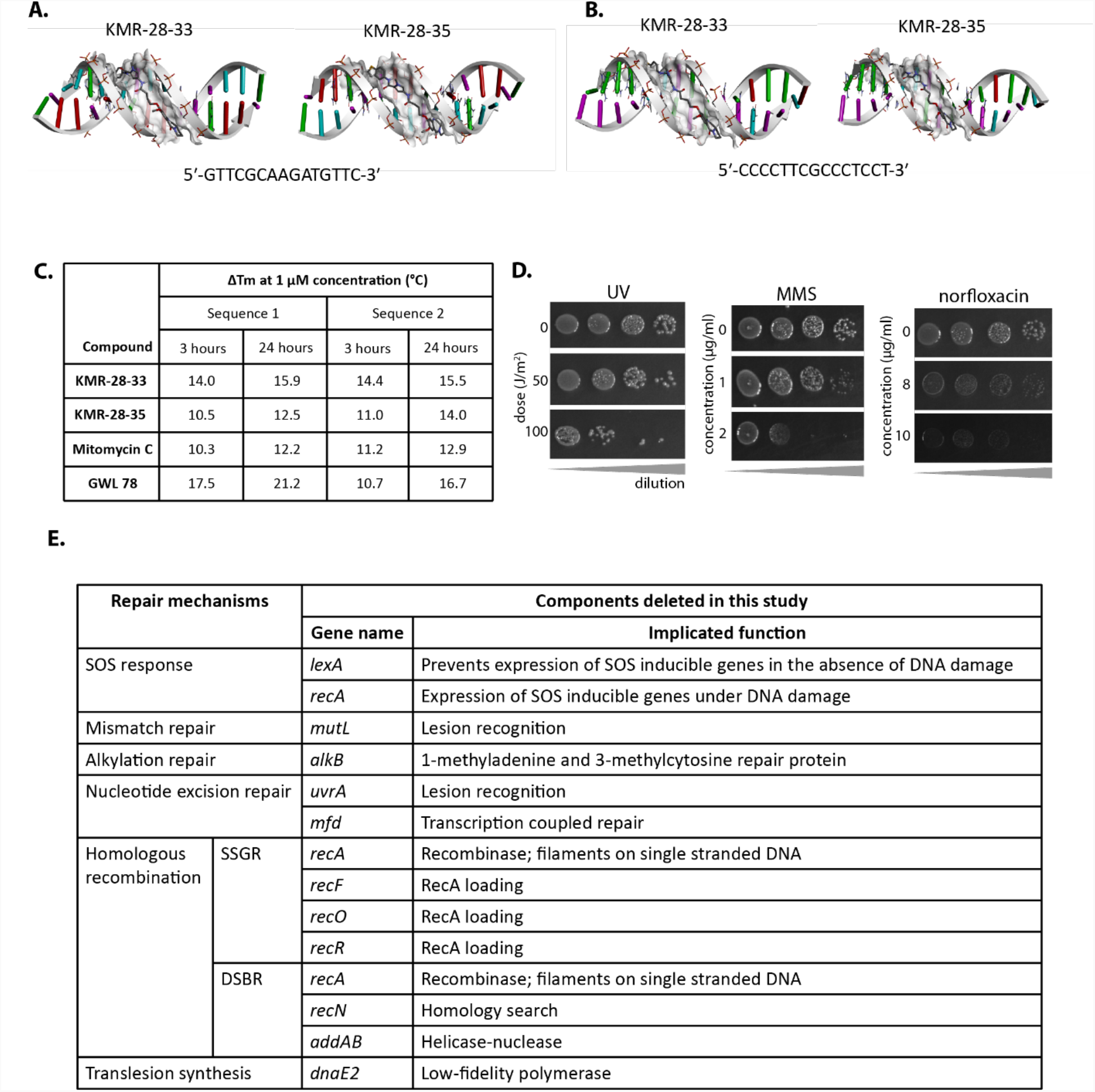
(A) Molecular docking of KMR-28-33 and KMR-28-35 with 15 bp DNA sequence taken from the LexA box within the promoter of *recA* gene (5’-GTTCGCAAGATGTTC-3’; GC content – 46%). (B) Molecular docking of KMR-28-33 and KMR-28-35 with 15 bp DNA sequence taken from the sRNA gene *CCNA_R0074* (5’-CCCCTTCGCCCTCCT-3’; GC content – 73%) (C) DNA thermal stabilization data for KMR-28-33, KMR-28-35 and mitomycin C with an AT-rich (seq-1) and a GC-rich (seq-2) hairpin DNA sequence. As a positive control, previously characterized compound GWL-78 is used. Average from three replicates shown (see methods for details of experimental setup). (D) Representative images of wild type *Caulobacter crescentus* growth on increasing concentrations of DNA damaging agents UV, MMS and norfloxacin. Grey triangle at the bottom of each image panel depicts increasing dilution of the bacterial culture from left to right. Minimum of three independent experiments were performed for each condition. (E) Table of all the repair components deleted in this study and their ascribed functions; SSGR – single-strand gap repair, DSBR – double-strand break repair.

**Figure S3:**
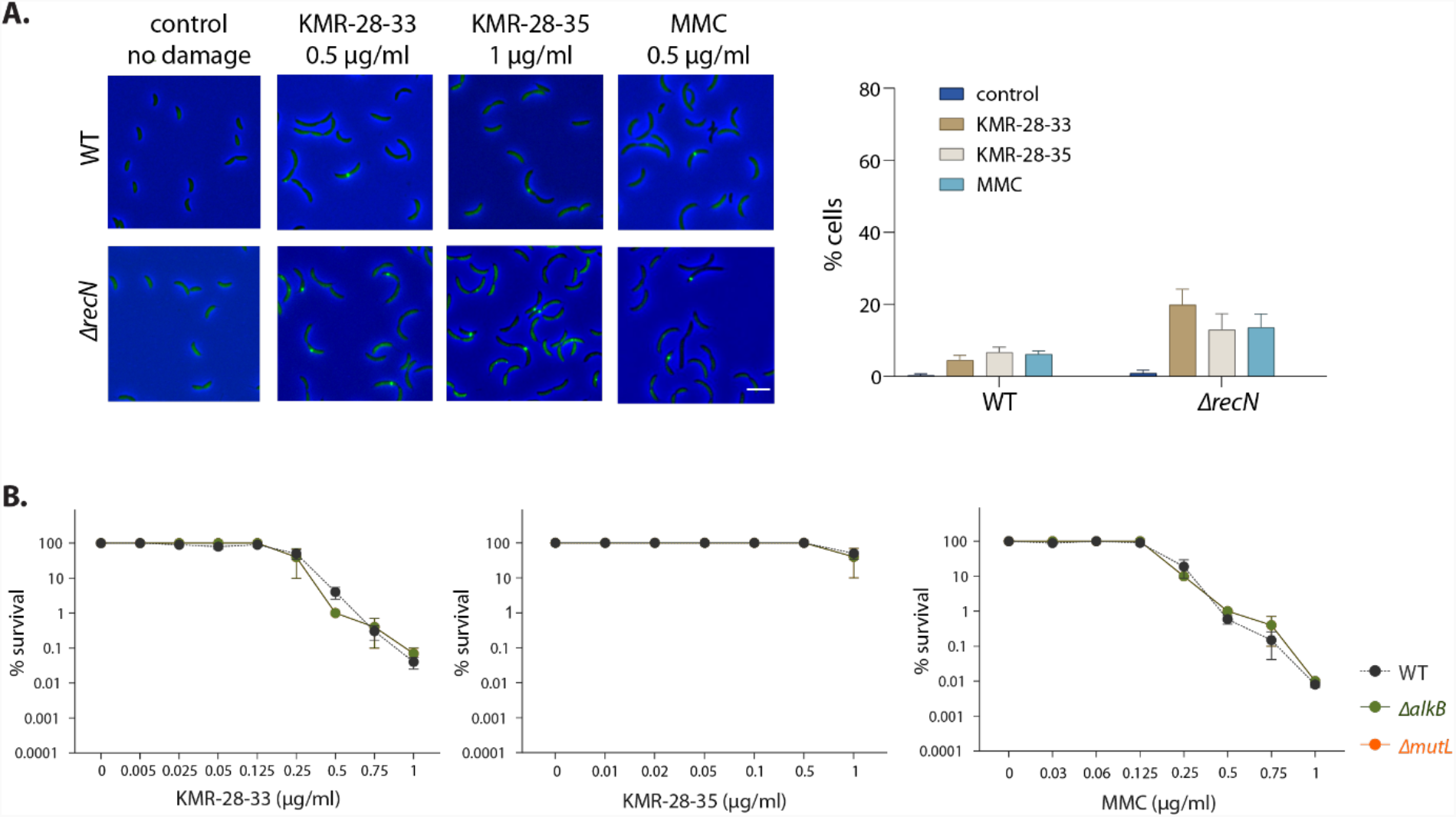
(A) [left] Representative images for cells with Gam-GFP foci upon treatment with KMR-28-33, KMR-28-35 and MMC in wild type and *ΔrecN* strains. Scale bar – 4 µm. [right] Percentage cells with Gam-GFP foci upon treatment with KMR-28-33, KMR-28-35 and MMC in wild type and *ΔrecN* strains. Mean and SD for data from three independent experiments is plotted (n *≤* 330 cells). (B) Survival of wild type, *ΔmutL* and *ΔalkB* strains under increasing doses of KMR-28-33, KMR-28-35 and MMC. Minimum of three independent experiments were performed for each strain. Mean and SEM from all repeats for each strain is plotted (wild type data from Figure 3A for comparison).

**Figure S4:**
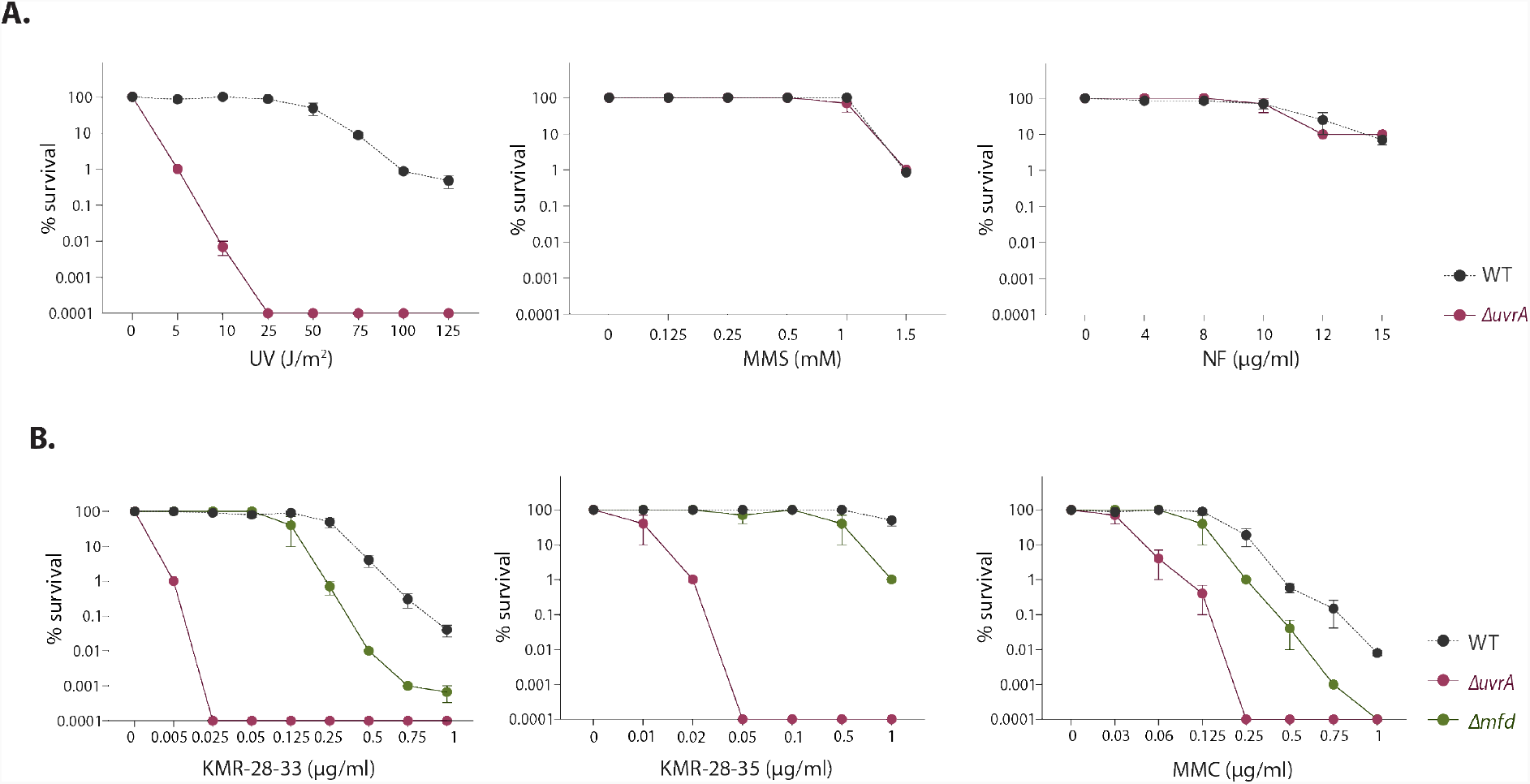
(A) Survival of wild type and *ΔuvrA* strains under increasing doses of UV, MMS and norfloxacin (NF) damage. Minimum of three independent experiments were performed for each strain. Mean and SEM from all repeats for each strain is plotted. (B) Survival of wild type, *ΔuvrA* and *Δmfd* strains under increasing doses of KMR-28-33, KMR-28-35 and MMC. Minimum of three independent experiments were performed for each strain. Mean and SEM from all repeats for each strain is plotted (wild type and *ΔuvrA* data from Figure 3A for comparison).

**Figure S5:**
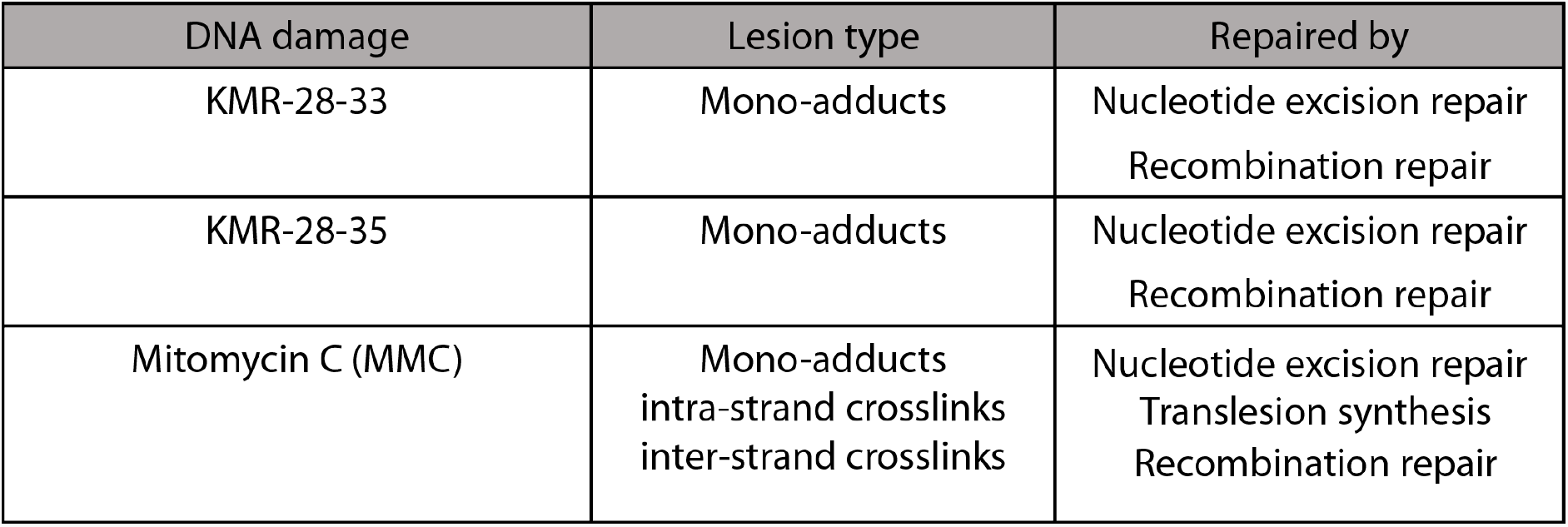
Table summarizing findings from this study on DNA repair mechanisms essential for repair/tolerance of lesions induced by KMR-28-33, KMR-28-35 and MMC.

**Table S1:** Strains used in present study.

| Strain name | Genotype | Strain construction |
| --- | --- | --- |
| CB15N |  |  |
| NABC2 | *CB15N; ΔrecA* | (Modell et al., 2014) |
| NABC29 | *CB15N; ΔdnaE2* | CB15N was transformed with pNABC148 plasmid to generate deletion of *dnaE2* through two-step recombination procedure. |
| NABC238 | *CB15N; ΔuvrA* | CB15N transformed with pNABC417 plasmid to generate deletion of *uvrA* through two-step recombination procedure. |
| NABC239 | *CB15N; ΔaddAB::gent* | CB15N transduced with lysate of strain harboring deletion of *addAB* linked to *gent<sup>R</sup>* |
| NABC265 | *CB15N; ΔrecA::kan<sup>R</sup>, ΔlexA::tet<sup>R</sup>, ΔsidA* | *CB15N; ΔlexA::tet<sup>R</sup>, ΔsidA* strain transduced with lysate of strain harboring *recA* deletion linked to *kan<sup>R</sup>* |
| NABC268 | *CB15N; P<sub>xyI</sub>-Gam-GFP::spec<sup>R</sup>* | CB15N transformed with pNABC592 plasmid |
| NABC439 | *CB15N; ΔrecN* | (Badrinarayanan et al., 2015) |
| NABC499 | *CB15N; ΔrecF* | CB15N transformed with pYY193 plasmid to generate deletion of *recF* (3rd-100<sup>th</sup> codon) through two-step recombination procedure. |
| NABC502 | *CB15N; ΔrecR* | CB15N transformed with pYY115 plasmid to generate deletion of *recR* through two-step recombination procedure. |
| NABC505 | *CB15N; ΔrecO* | CB15N transformed with pYY119 plasmid to generate deletion of *recO* through two-step recombination procedure. |
| NABC544 | *CB15N; Δmfd* | CB15N transformed with pNABC590 plasmid to generate deletion of *mfd* through two-step recombination procedure. |
| NABC579 | *CB15N; ΔalkB* | CB15N transformed with pNABC591 plasmid to generate deletion of *alkB* through two-step recombination procedure. |
| NABC580 | *CB15N; ΔmutL* | CB15N transformed with pNABC416 plasmid to generate deletion of *mutL* through two-step recombination procedure. |
| NABC581 | *CB15N; P<sub>sidA</sub>-YFP::kan<sup>R</sup>* | CB15N transformed with pNABC420 (Chimthanawala et al., 2021) |
| NABC582 | *CB15N; ΔuvrA; P<sub>xyI</sub>-Gam-GFP::spec<sup>R</sup>* | NABC238 transformed with pNABC592 plasmid |
| NABC583 | *CB15N; ΔrecN::spec<sup>R</sup>; P<sub>xyI</sub>-Gam-GFP::spec<sup>R</sup>* | NABC439 transformed with pNABC592 plasmid |

**Table S2:** Plasmids used in present study.

| Plasmid name | Construct details | Antibiotic marker |
| --- | --- | --- |
| pNPTS138 | (Skerker et al., 2005) | Kanamycin |
| pXGFPC1 | (Thanbichler et al., 2007) | Spectinomycin |
| pXYFPC2 | (Thanbichler et al., 2007) | Kanamycin |
| pNABC148 | 600 bp fragments upstream and downstream of *dnaE2* genomic locus were amplified from *C. crescentus* gDNA using RR_oligo_021/RR_oligo_022 (upstream fragment) and RR_oligo_023/RR_oligo_024 (downstream fragment) primer pairs. These fragments were assembled with BamH1/Nhe1 linearized pNPTS138 vector using Gibson assembly. | Kanamycin |
| pNABC416 | 600 bp fragments upstream and downstream of *mutL* genomic locus were amplified from *C. crescentus* gDNA using PS_oligo_049/AMJ_oligo_061 (upstream fragment) and AMJ_oligo_062/PS_oligo_054 (downstream fragment) primer pairs. These fragments were assembled with BamH1/Nhe1 linearized pNPTS138 vector using Gibson assembly. | Kanamycin |
| pNABC417 | 600 bp fragments upstream and downstream of *uvrA* genomic locus were amplified from *C. crescentus* gDNA using PS_oligo_037/AMJ_oligo_057 (upstream fragment) and AMJ_oligo_058/PS_oligo_042 (downstream fragment) primer pairs. These fragments were assembled with BamH1/Nhe1 linearized pNPTS138 vector using Gibson assembly. | Kanamycin |
| pYY193 | 500 bp fragments upstream and downstream of *recF* (from the 3<sup>rd</sup> to the 100<sup>th</sup> codon) were amplified from *C. crescentus* gDNA using UPdel00158_F/UPdel00158_R (upstream fragment) and DWNdel00158_F/DWNdel00158_R (downstream fragment) primer pairs. These fragments were assembled with EcoRI/BamHI linearized pNPTS138 vector using Gibson assembly. | Kanamycin |
| pYY115 | 500 bp fragments upstream and downstream of *recR* were amplified from *C. crescentus* gDNA using UPdel00270_F/UPdel00270_R (upstream fragment) and DWNdel00270_F/DWNdel00270_R (downstream fragment) primer pairs. These fragments were assembled with EcoRI/BamHI linearized pNPTS138 vector using Gibson assembly. | Kanamycin |
| pYY119 | 500 bp fragments upstream and downstream of *recO* were amplified from *C. crescentus* gDNA using UPdel01635_F/UPdel01635_R (upstream fragment) and DWNdel01635_F/DWNdel01635_R (downstream fragment) primer pairs. These fragments were assembled with EcoRI/BamHI linearized pNPTS138 vector using Gibson assembly. | Kanamycin |
| pNABC420 | pXYFPC2 vector was amplified using AB_oligo_651 and AB_oligo_652 and P<sub>sidA</sub>-YFP fragment was amplified from a replicating plasmid harbouring YFP under P<sub>sidA</sub> promoter using AC_oligo_322 and AC_oligo_321. The vector and insert fragments were assembled with Gibson assembly. | Kanamycin |
| pNABC590 | 600 bp fragments upstream and downstream of *mfd* genomic locus were amplified from *C. crescentus* gDNA using SD_oligo_088/SD_oligo_089 (upstream fragment) and SD_oligo_090/SD_oligo_0091 (downstream fragment) primer pairs. These fragments were assembled with BamHI/NheI linearized pNPTS138 vector using Gibson assembly. | Kanamycin |
| pNABC591 | 600 bp fragments upstream and downstream of *alkB* genomic locus were amplified from *C. crescentus* gDNA using AMJ_oligo_041/AMJ_oligo_053 (upstream fragment) and AMJ_oligo_054/PS_oligo_044 (downstream fragment) primer pairs. These fragments were assembled with BamHI/NheI linearized pNPTS138 vector using Gibson assembly. | Kanamycin |
| pNABC592 | *gam_GFP* was amplified using SD_oligo_019 and SD_oligo_020 from an *E. coli* strain harboring *gam-gfp* (Shee *et al.*, 2013) and assembled with NdeI/NheI digested pXYFPC-1 using Gibson assembly. | Spectinomycin |

**Table S3:**
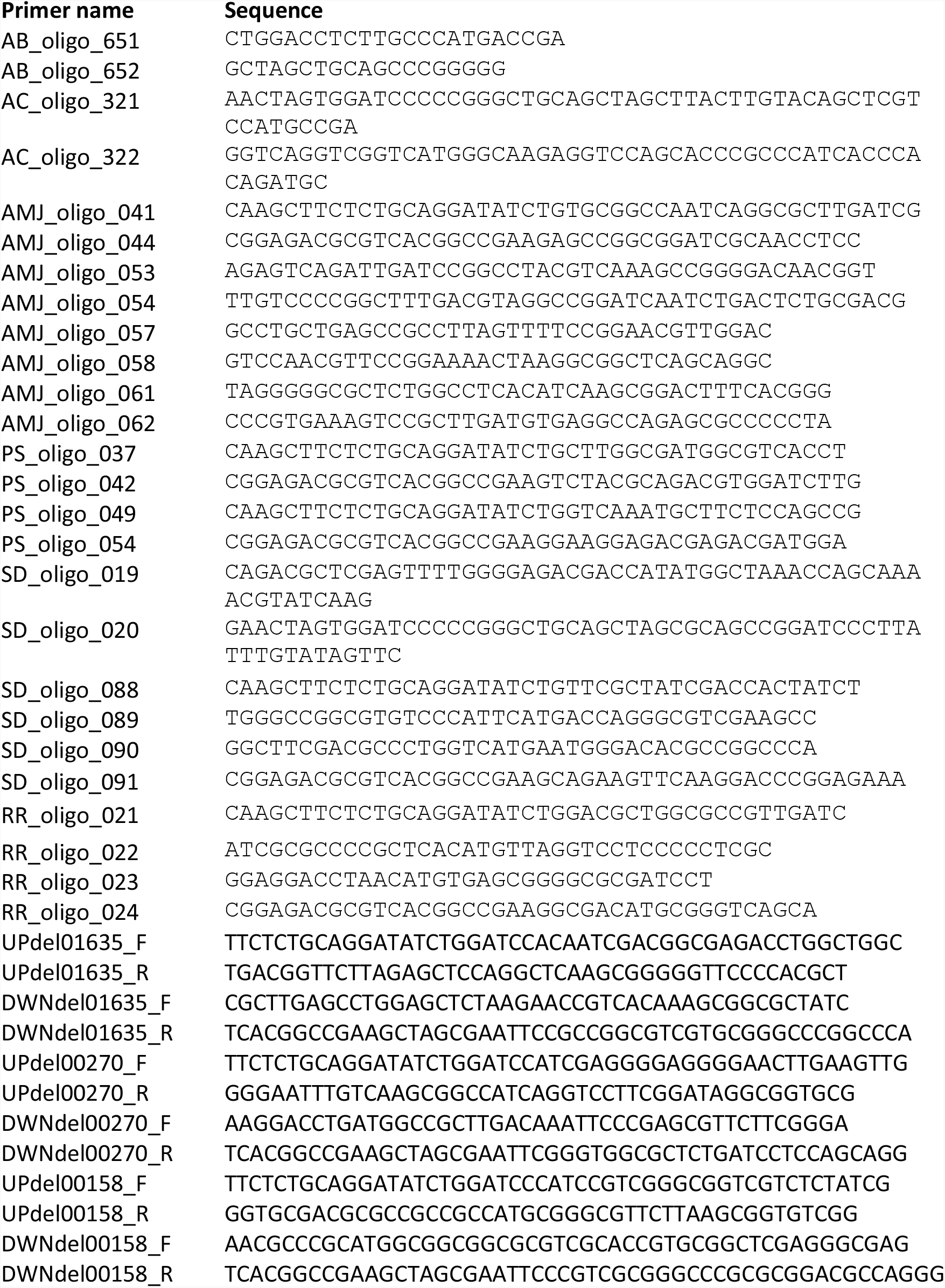
Oligos used in present study.

## References

Aguilera, A., & Gaillard, H. (2014). Transcription and Recombination: When RNA Meets DNA. Cold Spring Harbor Perspectives in Biology, 6(8), a016543–a016543. https://doi.org/10.1101/cshperspect.a016543

Alves, I. R., Lima-Noronha, M. A., Silva, L. G., Fernández-Silva, F. S., Freitas, A. L. D., Marques, M. V., & Galhardo, R. S. (2017). Effect of SOS-induced levels of imuABC on spontaneous and damage-induced mutagenesis in Caulobacter crescentus. DNA Repair, 59, 20–26. https://doi.org/10.1016/j.dnarep.2017.09.003

Amundsen, S. K., Spicer, T., Karabulut, A. C., Londoño, L. M., Eberhart, C., Fernandez Vega, V., Bannister, T. D., Hodder, P., & Smith, G. R. (2012). Small-Molecule Inhibitors of Bacterial AddAB and RecBCD Helicase-Nuclease DNA Repair Enzymes. ACS Chemical Biology, 7(5), 879–891. https://doi.org/10.1021/cb300018x

Andriollo, P., Hind, C. K., Picconi, P., Nahar, K. S., Jamshidi, S., Varsha, A., Clifford, M., Sutton, J. M., & Rahman, K. M. (2018). C8-Linked Pyrrolobenzodiazepine Monomers with Inverted Building Blocks Show Selective Activity against Multidrug Resistant Gram-Positive Bacteria. ACS Infectious Diseases, 4(2), 158–174. https://doi.org/10.1021/acsinfecdis.7b00130

Arnould, S., Spanswick, V. J., Macpherson, J. S., Hartley, J. A., Thurston, D. E., Jodrell, D. I., & Guichard, S. M. (2006). Time-dependent cytotoxicity induced by SJG-136 (NSC 694501): Influence of the rate of interstrand cross-link formation on DNA damage signaling. Molecular Cancer Therapeutics, 5(6), 1602–1609. https://doi.org/10.1158/1535-7163.MCT-06-0018

Badrinarayanan, A., Le, T. B. K., & Laub, M. T. (2015). Rapid pairing and resegregation of distant homologous loci enables double-strand break repair in bacteria. The Journal of Cell Biology, 210(3), 385–400. https://doi.org/10.1083/jcb.201505019

Bargonetti, J., Champeil, E., & Tomasz, M. (2010). Differential Toxicity of DNA Adducts of Mitomycin C. Journal of Nucleic Acids, 2010, 1–6. https://doi.org/10.4061/2010/698960

Beranek, D. T. (1990). Distribution of methyl and ethyl adducts following alkylation with monofunctional alkylating agents. Mutation Research/Fundamental and Molecular Mechanisms of Mutagenesis, 231(1), 11–30. https://doi.org/10.1016/0027-5107(90)90173-2

Bose, D. S., Thompson, A. S., Ching, J., Hartley, J. A., Berardini, M. D., Jenkins, T. C., Neidle, S., Hurley, L. H., & Thurston, D. E. (1992). Rational design of a highly efficient irreversible DNA interstrand cross-linking agent based on the pyrrolobenzodiazepine ring system. Journal of the American Chemical Society, 114(12), 4939–4941. https://doi.org/10.1021/ja00038a089

Boshoff, H. I. M., Reed, M. B., Barry, C. E., & Mizrahi, V. (2003). DnaE2 polymerase contributes to in vivo survival and the emergence of drug resistance in Mycobacterium tuberculosis. Cell, 113(2), 183–193.

Brucoli, F., Guzman, J. D., Basher, M. A., Evangelopoulos, D., McMahon, E., Munshi, T., McHugh, T. D., Fox, K. R., & Bhakta, S. (2016). DNA sequence-selective C8-linked pyrrolobenzodiazepine–heterocyclic polyamide conjugates show anti-tubercular-specific activities. The Journal of Antibiotics, 69(12), 843–849. https://doi.org/10.1038/ja.2016.43

Burby, P. E., & Simmons, L. A. (2019). A bacterial DNA repair pathway specific to a natural antibiotic. Molecular Microbiology, 111(2), 338–353. https://doi.org/10.1111/mmi.14158

Chai, T., Terrettaz, C., & Collier, J. (2021). Spatial coupling between DNA replication and mismatch repair in Caulobacter crescentus. Nucleic Acids Research, 49(6), 3308–3321. https://doi.org/10.1093/nar/gkab112

Chatterjee, N., & Walker, G. C. (2017). Mechanisms of DNA damage, repair, and mutagenesis. Environmental and Molecular Mutagenesis, 58(5), 235–263. https://doi.org/10.1002/em.22087

Cole, J. M., Acott, J. D., Courcelle, C. T., & Courcelle, J. (2018). Limited Capacity or Involvement of Excision Repair, Double-Strand Breaks, or Translesion Synthesis for Psoralen Cross-Link Repair in Escherichia coli. Genetics, 210(1), 99–112. https://doi.org/10.1534/genetics.118.301239

Colombi, D., & Gomes, S. L. (1997). An alkB gene homolog is differentially transcribed during the Caulobacter crescentus cell cycle. Journal of Bacteriology, 179(10), 3139–3145. https://doi.org/10.1128/jb.179.10.3139-3145.1997

Corcoran, D. B., Lewis, T., Nahar, K. S., Jamshidi, S., Fegan, C., Pepper, C., Thurston, D. E., & Rahman, K. Miraz. (2019). Effects of Systematic Shortening of Noncovalent C8 Side Chain on the Cytotoxicity and NF-κB Inhibitory Capacity of Pyrrolobenzodiazepines (PBDs). Journal of Medicinal Chemistry, 62(4), 2127–2139. https://doi.org/10.1021/acs.jmedchem.8b01849

de Almeida, L. C., Calil, F. A., Machado-Neto, J. A., & Costa-Lotufo, L. V. (2021). DNA damaging agents and DNA repair: From carcinogenesis to cancer therapy. Cancer Genetics, 252–253, 6–24. https://doi.org/10.1016/j.cancergen.2020.12.002

Ely, B. (1991). [17] Genetics of Caulobacter crescentus. In Methods in Enzymology (Vol. 204, pp. 372–384). Elsevier. https://doi.org/10.1016/0076-6879(91)04019-K

Fitzgerald, D. M., Hastings, P. J., & Rosenberg, S. M. (2017). Stress-Induced Mutagenesis: Implications in Cancer and Drug Resistance. Annual Review of Cancer Biology, 1(1), 119–140. https://doi.org/10.1146/annurev-cancerbio-050216-121919

Galhardo, R. S. (2005). An SOS-regulated operon involved in damage-inducible mutagenesis in Caulobacter crescentus. Nucleic Acids Research, 33(8), 2603–2614. https://doi.org/10.1093/nar/gki551

Gerratana, B. (2012). Biosynthesis, synthesis, and biological activities of pyrrolobenzodiazepines: ACTIVITIES OF PYRROLOBENZODIAZEPINES. Medicinal Research Reviews, 32(2), 254–293. https://doi.org/10.1002/med.20212

Gregson, S. J., Howard, P. W., Hartley, J. A., Brooks, N. A., Adams, L. J., Jenkins, T. C., Kelland, L. R., & Thurston, D. E. (2001). Design, Synthesis, and Evaluation of a Novel Pyrrolobenzodiazepine DNA-Interactive Agent with Highly Efficient Cross-Linking Ability and Potent Cytotoxicity. Journal of Medicinal Chemistry, 44(5), 737–748. https://doi.org/10.1021/jm001064n

Hartley, J. A., Hamaguchi, A., Coffils, M., Martin, C. R. H., Suggitt, M., Chen, Z., Gregson, S. J., Masterson, L. A., Tiberghien, A. C., Hartley, J. M., Pepper, C., Lin, T. T., Fegan, C., Thurston, D. E., & Howard, P. W. (2010). SG2285, a Novel C2-Aryl-Substituted Pyrrolobenzodiazepine Dimer Prodrug That Cross-links DNA and Exerts Highly Potent Antitumor Activity. Cancer Research, 70(17), 6849–6858. https://doi.org/10.1158/0008-5472.CAN-10-0790

Hoffmann, R. M., Crescioli, S., Mele, S., Sachouli, E., Cheung, A., Chui, C. K., Andriollo, P., Jackson, P. J. M., Lacy, K. E., Spicer, J. F., Thurston, D. E., & Karagiannis, S. N. (2020). A Novel Antibody-Drug Conjugate (ADC) Delivering a DNA Mono-Alkylating Payload to Chondroitin Sulfate Proteoglycan (CSPG4)-Expressing Melanoma. Cancers, 12(4), E1029. https://doi.org/10.3390/cancers12041029

Hurley, L. H. (1977). Pyrrolo(1,4)benzodiazepine antitumor antibiotics. Comparative aspects of anthramycin, tomaymycin and sibiromycin. The Journal of Antibiotics, 30(5), 349–370. https://doi.org/10.7164/antibiotics.30.349

Inomata, Y., Abe, T., Tsuda, M., Takeda, S., & Hirota, K. (2021). Division of labor of Y-family polymerases in translesion-DNA synthesis for distinct types of DNA damage. PLOS ONE, 16(6), e0252587. https://doi.org/10.1371/journal.pone.0252587

Ippoliti, P. J., DeLateur, N. A., Jones, K. M., & Beuning, P. J. (2012). Multiple Strategies for Translesion Synthesis in Bacteria. Cells, 1(4), 799–831. https://doi.org/10.3390/cells1040799

Jackson, P. J. M., Kay, S., Pysz, I., & Thurston, D. E. (2018). Use of pyrrolobenzodiazepines and related covalent-binding DNA-interactive molecules as ADC payloads: Is mechanism related to systemic toxicity? Drug Discovery Today: Technologies, 30, 71–83. https://doi.org/10.1016/j.ddtec.2018.10.004

Jatsenko, T., Sidorenko, J., Saumaa, S., & Kivisaar, M. (2017). DNA Polymerases ImuC and DinB Are Involved in DNA Alkylation Damage Tolerance in Pseudomonas aeruginosa and Pseudomonas putida. PLOS ONE, 12(1), e0170719. https://doi.org/10.1371/journal.pone.0170719

Jenkins, T. C., Hurley, L. H., Neidle, S., & Thurston, D. E. (1994). Structure of a Covalent DNA Minor Groove Adduct with a Pyrrolobenzodiazepine Dimer: Evidence for Sequence-Specific Interstrand Crosslinking. Journal of Medicinal Chemistry, 37(26), 4529–4537. https://doi.org/10.1021/jm00052a012

Jia, L., Kropachev, K., Ding, S., Van Houten, B., Geacintov, N. E., & Broyde, S. (2009). Exploring damage recognition models in prokaryotic nucleotide excision repair with a benzo[a]pyrene-derived lesion in UvrB. Biochemistry, 48(38), 8948–8957. https://doi.org/10.1021/bi9010072

Joseph, A. M., & Badrinarayanan, A. (2020). Visualizing mutagenic repair: Novel insights into bacterial translesion synthesis. FEMS Microbiology Reviews, 44(5), 572–582. https://doi.org/10.1093/femsre/fuaa023

Joseph, A. M., Daw, S., Sadhir, I., & Badrinarayanan, A. (2021). Coordination between nucleotide excision repair and specialized polymerase DnaE2 action enables DNA damage survival in non-replicating bacteria [Preprint]. Microbiology. https://doi.org/10.1101/2021.02.15.431208

Kisker, C., Kuper, J., & Van Houten, B. (2013). Prokaryotic nucleotide excision repair. Cold Spring Harbor Perspectives in Biology, 5(3), a012591. https://doi.org/10.1101/cshperspect.a012591

Kotecha, M., Kluza, J., Wells, G., O’Hare, C. C., Forni, C., Mantovani, R., Howard, P. W., Morris, P., Thurston, D. E., Hartley, J. A., & Hochhauser, D. (2008). Inhibition of DNA binding of the NF-Y transcription factor by the pyrrolobenzodiazepine-polyamide conjugate GWL-78. Molecular Cancer Therapeutics, 7(5), 1319–1328. https://doi.org/10.1158/1535-7163.MCT-07-0475

Kovtun, Y., Noordhuis, P., Whiteman, K. R., Watkins, K., Jones, G. E., Harvey, L., Lai, K. C., Portwood, S., Adams, S., Sloss, C. M., Schuurhuis, G. J., Ossenkoppele, G., Wang, E. S., & Pinkas, J. (2018). IMGN779, a Novel CD33-Targeting Antibody–Drug Conjugate with DNA-Alkylating Activity, Exhibits Potent Antitumor Activity in Models of AML. Molecular Cancer Therapeutics, 17(6), 1271–1279. https://doi.org/10.1158/1535-7163.MCT-17-1077

Kung Sutherland, M. S., Walter, R. B., Jeffrey, S. C., Burke, P. J., Yu, C., Kostner, H., Stone, I., Ryan, M. C., Sussman, D., Lyon, R. P., Zeng, W., Harrington, K. H., Klussman, K., Westendorf, L., Meyer, D., Bernstein, I. D., Senter, P. D., Benjamin, D. R., Drachman, J. G., & McEarchern, J. A. (2013). SGN-CD33A: A novel CD33-targeting antibody–drug conjugate using a pyrrolobenzodiazepine dimer is active in models of drug-resistant AML. Blood, 122(8), 1455–1463. https://doi.org/10.1182/blood-2013-03-491506

Leimgruber, W., Stefanović, V., Schenker, F., Karr, A., & Berger, J. (1965). Isolation and Characterization of Anthramycin, a New Antitumor Antibiotic. Journal of the American Chemical Society, 87(24), 5791–5793. https://doi.org/10.1021/ja00952a050

Lim, C. S. Q., Ha, K. P., Clarke, R. S., Gavin, L.-A., Cook, D. T., Hutton, J. A., Sutherell, C. L., Edwards, A. M., Evans, L. E., Tate, E. W., & Lanyon-Hogg, T. (2019). Identification of a potent small-molecule inhibitor of bacterial DNA repair that potentiates quinolone antibiotic activity in methicillin-resistant Staphylococcus aureus. Bioorganic & Medicinal Chemistry, 27(20), 114962. https://doi.org/10.1016/j.bmc.2019.06.025

Liu, Y., Reeves, D., Kropachev, K., Cai, Y., Ding, S., Kolbanovskiy, M., Kolbanovskiy, A., Bolton, J. L., Broyde, S., Van Houten, B., & Geacintov, N. E. (2011). Probing for DNA damage with β-hairpins: Similarities in incision efficiencies of bulky DNA adducts by prokaryotic and human nucleotide excision repair systems in vitro. DNA Repair, 10(7), 684–696. https://doi.org/10.1016/j.dnarep.2011.04.020

Lopes-Kulishev, C. O., Alves, I. R., Valencia, E. Y., Pidhirnyj, M. I., Fernández-Silva, F. S., Rodrigues, T. R., Guzzo, C. R., & Galhardo, R. S. (2015). Functional characterization of two SOS-regulated genes involved in mitomycin C resistance in Caulobacter crescentus. DNA Repair, 33, 78–89. https://doi.org/10.1016/j.dnarep.2015.06.009

Mantaj, J., Jackson, P. J. M., Rahman, K. M., & Thurston, D. E. (2017). From Anthramycin to Pyrrolobenzodiazepine (PBD)-Containing Antibody-Drug Conjugates (ADCs). Angewandte Chemie International Edition, 56(2), 462–488. https://doi.org/10.1002/anie.201510610

Mazloum, N., Stegman, M. A., Croteau, D. L., Van Houten, B., Kwon, N. S., Ling, Y., Dickinson, C., Venugopal, A., Towheed, M. A., & Nathan, C. (2011). Identification of a Chemical That Inhibits the Mycobacterial UvrABC Complex in Nucleotide Excision Repair. Biochemistry, 50(8), 1329–1335. https://doi.org/10.1021/bi101674c

Mehta, A., & Haber, J. E. (2014). Sources of DNA Double-Strand Breaks and Models of Recombinational DNA Repair. Cold Spring Harbor Perspectives in Biology, 6(9), a016428–a016428. https://doi.org/10.1101/cshperspect.a016428

Modell, J. W., Hopkins, A. C., & Laub, M. T. (2011). A DNA damage checkpoint in Caulobacter crescentus inhibits cell division through a direct interaction with FtsW. Genes & Development, 25(12), 1328–1343. https://doi.org/10.1101/gad.2038911

Morgensztern, D., Besse, B., Greillier, L., Santana-Davila, R., Ready, N., Hann, C. L., Glisson, B. S., Farago, A. F., Dowlati, A., Rudin, C. M., Le Moulec, S., Lally, S., Yalamanchili, S., Wolf, J., Govindan, R., & Carbone, D. P. (2019). Efficacy and Safety of Rovalpituzumab Tesirine in Third-Line and Beyond Patients with DLL3-Expressing, Relapsed/Refractory Small-Cell Lung Cancer: Results From the Phase II TRINITY Study. Clinical Cancer Research, 25(23), 6958–6966. https://doi.org/10.1158/1078-0432.CCR-19-1133

Ona, K. R., Courcelle, C. T., & Courcelle, J. (2009). Nucleotide Excision Repair Is a Predominant Mechanism for Processing Nitrofurazone-Induced DNA Damage in Escherichia coli. Journal of Bacteriology, 191(15), 4959–4965. https://doi.org/10.1128/JB.00495-09

Pages, V. (2003). Uncoupling of Leading-and Lagging-Strand DNA Replication During Lesion Bypass in Vivo. Science, 300(5623), 1300–1303. https://doi.org/10.1126/science.1083964

Paintdakhi, A., Parry, B., Campos, M., Irnov, I., Elf, J., Surovtsev, I., & Jacobs-Wagner, C. (2016). Oufti: An integrated software package for high-accuracy, high-throughput quantitative microscopy analysis. Molecular Microbiology, 99(4), 767–777. https://doi.org/10.1111/mmi.13264

Picconi, P., Hind, C. K., Nahar, K. S., Jamshidi, S., Di Maggio, L., Saeed, N., Evans, B., Solomons, J., Wand, M. E., Sutton, J. M., & Rahman, K. M. (2020). New Broad-Spectrum Antibiotics Containing a Pyrrolobenzodiazepine Ring with Activity against Multidrug-Resistant Gram-Negative Bacteria. Journal of Medicinal Chemistry, 63(13), 6941–6958. https://doi.org/10.1021/acs.jmedchem.0c00328

Prakash, S., Johnson, R. E., & Prakash, L. (2005). EUKARYOTIC TRANSLESION SYNTHESIS DNA POLYMERASES: Specificity of Structure and Function. Annual Review of Biochemistry, 74(1), 317–353. https://doi.org/10.1146/annurev.biochem.74.082803.133250

Puzanov, I., Lee, W., Chen, A. P., Calcutt, M. W., Hachey, D. L., Vermeulen, W. L., Spanswick, V. J., Liao, C.-Y., Hartley, J. A., Berlin, J. D., & Rothenberg, M. L. (2011). Phase I Pharmacokinetic and Pharmacodynamic Study of SJG-136, a Novel DNA Sequence Selective Minor Groove Cross-linking Agent, in Advanced Solid Tumors. Clinical Cancer Research, 17(11), 3794–3802. https://doi.org/10.1158/1078-0432.CCR-10-2056

Rahman, K. M., Jackson, P. J. M., James, C. H., Basu, B. P., Hartley, J. A., de la Fuente, M., Schatzlein, A., Robson, M., Pedley, R. B., Pepper, C., Fox, K. R., Howard, P. W., & Thurston, D. E. (2013). GC-Targeted C8-Linked Pyrrolobenzodiazepine–Biaryl Conjugates with Femtomolar in Vitro Cytotoxicity and in Vivo Antitumor Activity in Mouse Models. Journal of Medicinal Chemistry, 56(7), 2911–2935. https://doi.org/10.1021/jm301882a

Rahman, K. M., James, C. H., & Thurston, D. E. (2011). Effect of base sequence on the DNA cross-linking properties of pyrrolobenzodiazepine (PBD) dimers. Nucleic Acids Research, 39(13), 5800–5812. https://doi.org/10.1093/nar/gkr122

Rahman, K. M., Rosado, H., Moreira, J. B., Feuerbaum, E.-A., Fox, K. R., Stecher, E., Howard, P. W., Gregson, S. J., James, C. H., de la Fuente, M., Waldron, D. E., Thurston, D. E., & Taylor, P. W. (2012). Antistaphylococcal activity of DNA-interactive pyrrolobenzodiazepine (PBD) dimers and PBD-biaryl conjugates. Journal of Antimicrobial Chemotherapy, 67(7), 1683–1696. https://doi.org/10.1093/jac/dks127

Rocha, E. P. C., Cornet, E., & Michel, B. (2005). Comparative and Evolutionary Analysis of the Bacterial Homologous Recombination Systems. PLoS Genetics, 1(2), e15. https://doi.org/10.1371/journal.pgen.0010015

Rosado, H., Rahman, K. M., Feuerbaum, E.-A., Hinds, J., Thurston, D. E., & Taylor, P. W. (2011). The minor groove-binding agent ELB-21 forms multiple interstrand and intrastrand covalent cross-links with duplex DNA and displays potent bactericidal activity against methicillin-resistant Staphylococcus aureus. Journal of Antimicrobial Chemotherapy, 66(5), 985–996. https://doi.org/10.1093/jac/dkr044

Selby, C. P. (2017). Mfd Protein and Transcription-Repair Coupling in Escherichia coli. Photochemistry and Photobiology, 93(1), 280–295. https://doi.org/10.1111/php.12675

Selby, C., & Sancar, A. (1993). Molecular mechanism of transcription-repair coupling. Science, 260(5104), 53–58. https://doi.org/10.1126/science.8465200

Shee, C., Cox, B. D., Gu, F., Luengas, E. M., Joshi, M. C., Chiu, L.-Y., Magnan, D., Halliday, J. A., Frisch, R. L., Gibson, J. L., Nehring, R. B., Do, H. G., Hernandez, M., Li, L., Herman, C., Hastings, P., Bates, D., Harris, R. S., Miller, K. M., & Rosenberg, S. M. (2013). Engineered proteins detect spontaneous DNA breakage in human and bacterial cells. ELife, 2, e01222. https://doi.org/10.7554/eLife.01222

Skerker, J. M., Prasol, M. S., Perchuk, B. S., Biondi, E. G., & Laub, M. T. (2005). Two-Component Signal Transduction Pathways Regulating Growth and Cell Cycle Progression in a Bacterium: A System-Level Analysis. PLoS Biology, 3(10), e334. https://doi.org/10.1371/journal.pbio.0030334

Spies, M., & Kowalczykowski, S. C. (2014). Homologous Recombination by the RecBCD and RecF Pathways. In N. P. Higgins (Ed.), The Bacterial Chromosome (pp. 389–403). ASM Press. https://doi.org/10.1128/9781555817640.ch21

Strick, T. R., & Portman, J. R. (2019). Transcription-Coupled Repair: From Cells to Single Molecules and Back Again. Journal of Molecular Biology, 431(20), 4093–4102. https://doi.org/10.1016/j.jmb.2019.05.040

Surova, O., & Zhivotovsky, B. (2013). Various modes of cell death induced by DNA damage. Oncogene, 32(33), 3789–3797. https://doi.org/10.1038/onc.2012.556

Thanbichler, M., Iniesta, A. A., & Shapiro, L. (2007). A comprehensive set of plasmids for vanillate-and xylose-inducible gene expression in Caulobacter crescentus. Nucleic Acids Research, 35(20), e137. https://doi.org/10.1093/nar/gkm818

Thurston, D. E., Bose, D. S., Howard, P. W., Jenkins, T. C., Leoni, A., Baraldi, P. G., Guiotto, A., Cacciari, B., Kelland, L. R., Foloppe, M.-P., & Rault, S. (1999). Effect of A-Ring Modifications on the DNA-Binding Behavior and Cytotoxicity of Pyrrolo[2,1-c ][1,4]benzodiazepines. Journal of Medicinal Chemistry, 42(11), 1951–1964. https://doi.org/10.1021/jm981117p

Tomasz, M. (1995). Mitomycin C: Small, fast and deadly (but very selective). Chemistry & Biology, 2(9), 575–579. https://doi.org/10.1016/1074-5521(95)90120-5

Trott, O., & Olson, A. J. (2009). AutoDock Vina: Improving the speed and accuracy of docking with a new scoring function, efficient optimization, and multithreading. Journal of Computational Chemistry, NA-NA. https://doi.org/10.1002/jcc.21334

Warner, D. F., Ndwandwe, D. E., Abrahams, G. L., Kana, B. D., Machowski, E. E., Venclovas, C., & Mizrahi, V. (2010). Essential roles for imuA’-and imuB-encoded accessory factors in DnaE2-dependent mutagenesis in Mycobacterium tuberculosis. Proceedings of the National Academy of Sciences of the United States of America, 107(29), 13093–13098. https://doi.org/10.1073/pnas.1002614107

Warren, A. J., Maccubbin, A. E., & Hamilton, J. W. (1998). Detection of mitomycin C-DNA adducts in vivo by 32P-postlabeling: Time course for formation and removal of adducts and biochemical modulation. Cancer Research, 58(3), 453–461.

Waters, L. S., Minesinger, B. K., Wiltrout, M. E., D’Souza, S., Woodruff, R. V., & Walker, G. C. (2009). Eukaryotic Translesion Polymerases and Their Roles and Regulation in DNA Damage Tolerance. Microbiology and Molecular Biology Reviews, 73(1), 134–154. https://doi.org/10.1128/MMBR.00034-08

Wells, G., Martin, C. R. H., Howard, P. W., Sands, Z. A., Laughton, C. A., Tiberghien, A., Woo, C. K., Masterson, L. A., Stephenson, M. J., Hartley, J. A., Jenkins, T. C., Shnyder, S. D., Loadman, P. M., Waring, M. J., & Thurston, D. E. (2006). Design, Synthesis, and Biophysical and Biological Evaluation of a Series of Pyrrolobenzodiazepine-Poly(N -methylpyrrole) Conjugates. Journal of Medicinal Chemistry, 49(18), 5442–5461. https://doi.org/10.1021/jm051199z

Williams, H. L., Gottesman, M. E., & Gautier, J. (2013). The differences between ICL repair during and outside of S phase. Trends in Biochemical Sciences, 38(8), 386–393. https://doi.org/10.1016/j.tibs.2013.05.004

Xing, L., Lin, L., Yu, T., Li, Y., Wen, K., Cho, S.-F., Hsieh, P. A., Kinneer, K., Munshi, N. C., Anderson, K. C., & Tai, Y.-T. (2019). Anti-Bcma PBD MEDI2228 Combats Drug Resistance and Synergizes with Bortezomib and Inhibitors to DNA Damage Response in Multiple Myeloma. Blood, 134(Supplement_1), 1817–1817. https://doi.org/10.1182/blood-2019-127163

Zhong, H., Chen, C., Tammali, R., Breen, S., Zhang, J., Fazenbaker, C., Kennedy, M., Conway, J., Higgs, B. W., Holoweckyj, N., Raja, R., Harper, J., Pierce, A. J., Herbst, R., & Tice, D. A. (2019). Improved Therapeutic Window in BRCA -mutant Tumors with Antibody-linked Pyrrolobenzodiazepine Dimers with and without PARP Inhibition. Molecular Cancer Therapeutics, 18(1), 89–99. https://doi.org/10.1158/1535-7163.MCT-18-0314

## References

Chimthanawala, A., Parmar, J. J., Kumar, S., Iyer, K. S., Rao, M., & Badrinarayanan, A. (2021). SMC protein RecN drives RecA filament translocation and remodelling for in vivo homology search [Preprint]. Cell Biology. https://doi.org/10.1101/2021.08.16.456443

Modell, J. W., Kambara, T. K., Perchuk, B. S., & Laub, M. T. (2014). A DNA damage-induced, SOS-independent checkpoint regulates cell division in Caulobacter crescentus. PLoS Biology, 12(10), e1001977. https://doi.org/10.1371/journal.pbio.1001977

